# Dynamic perceptual feature selectivity in primary somatosensory cortex upon reversal learning

**DOI:** 10.1101/782847

**Authors:** Ronan Chéreau, Tanika Bawa, Leon Fodoulian, Alan Carleton, Stéphane Pagès, Anthony Holtmaat

## Abstract

Neurons in primary sensory cortex encode a variety of stimulus features upon perceptual learning. However, it is unclear whether the acquired stimulus selectivity remains stable when the same input is perceived in a different context. Here, we monitored the activity of individual neurons in the mouse primary somatosensory cortex in a reward-based texture discrimination task. We tracked their stimulus selectivity before and after changing reward contingencies, which allowed us to identify various classes of neurons. We found neurons that stably represented a texture or the upcoming behavioral choice, but the majority was dynamic. Among those, a subpopulation of neurons regained selectivity contingent on stimulus-value. These value-sensitive neurons forecasted the onset of learning by displaying a distinct and transient increase in activity, depending on past behavioral experience. Thus, stimulus selectivity of excitatory neurons during perceptual learning is dynamic and largely relies on behavioral contingencies, even in primary sensory cortex.

## INTRODUCTION

The mammalian cortex encodes a myriad of sensory signal characteristics which are represented by neuronal assemblies, each with a preference for specific stimulus parameters^1, 2^. It is believed that these assemblies are organized in a hierarchical fashion. First-order sensory areas encode lower-order stimulus features, such as textures coarseness^3,4,5^, object orientation and direction^6^, and sound frequency^7^, whereas more complex features and contextual aspects of a stimulus are encoded by higher-order cortices^8, 9, 10, 11^. Nonetheless, the coding in primary sensory cortices can exhibit higher levels of complexity, expressing non-sensory-related signals such as attention^12^, anticipation^13^ and behavioral choice^11, 12, 13, 14, 15^. Reward-based perceptual learning initially shapes the stimulus selectivity and response properties of primary sensory neurons, which may contribute to a reliable detection of particular features, and thereby improve perception^13, 16, 17, 18^. However, it is unclear as to whether the stimulus preference of those neurons remains stable when the reward contingencies are changed. To study this, we monitored the shaping of stimulus selectivity for primary somatosensory cortical (S1) layer 2/3 (L2/3) neurons in mice that learned to discriminate between a rewarded and non-rewarded texture. We then reassessed their selectivity upon reversal learning.

## RESULTS

### L2/3 neurons increase their selectivity for the rewarded texture with learning

We trained mice on a head-fixed ‘Go/No-go’ texture discrimination task, similar to previous designs^5^ (Fig. 1a). Thirsted animals were incited to lick a spout during a 2-second (s) presentation of a P120 sandpaper (Go stimulus), in order to trigger the supply of a water reward at the end of the texture presentation (‘hit’ trials) (Fig. 1a and S1). The animals needed to withhold from licking upon presentation of a P280 sandpaper (No-go stimulus) to avoid a 200-ms white noise and a 5-s timeout period. A failure to withhold from licking was scored as a ‘false alarm’ (FA) trial (Fig. 1a and Supplementary Fig. 1). Mice learned to discriminate between the two stimuli (Fig. 1b). They typically started at chance level (naïve) and reached an average of 82% performance level within 3-7 days, well above the expert criterion of 70% (Fig. 1b)^13, 14, 15, 19^. Trimming of the whiskers ipsilateral to the texture reduced the performance to chance level (Fig. 1c), indicating that the mice had fully relied on somatosensory input and had not used additional sensory information to solve the task. To confirm that S1 is required for this task, we suppressed its activity using a local injection of the γ-aminobutyric acid receptor (GABAR) agonist muscimol. This significantly decreased the performance of mice that had initially learned the task (Fig. 1d).

**Fig. 1.**
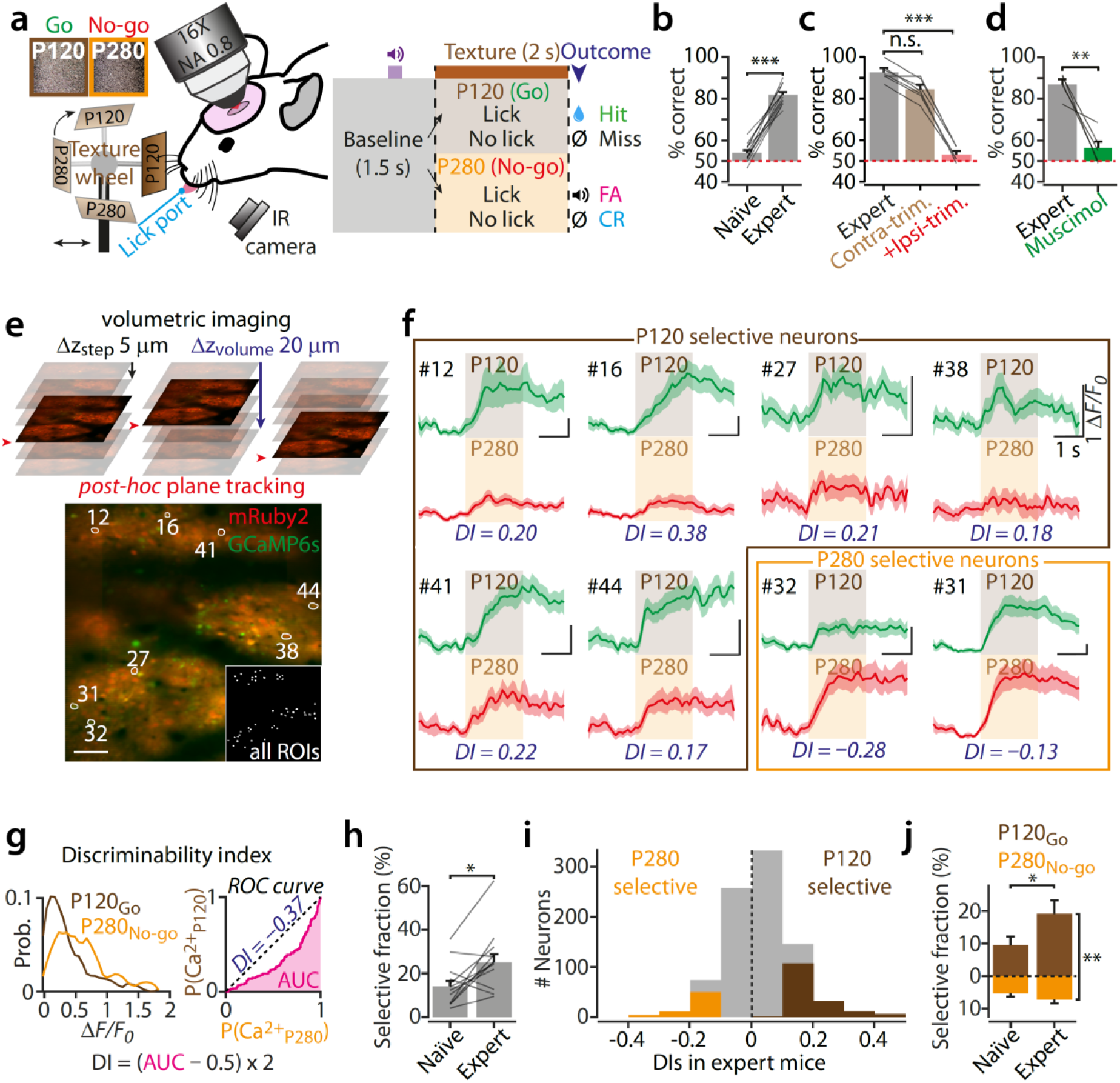
Increase in texture selectivity of S1 neurons during discrimination learning. **a**-Behavioral setup and experimental time-course. Mice were rewarded for licking the water spout upon random presentations of a P120 (Go stimulus) but not a P280 (No-go stimulus) sandpaper. **b**-Discrimination performance across learning (*N*=12 mice, Wilcoxon rank sum test, fraction of correct trials: naïve, 54.0 ± 1.2% (mean ± s.e.m.), expert, 81.8 ± 1.3%, \*\*\**P*<10^-8^). **c**-Effect of whisker trimming on behavioral performance (*N*=7 mice, Friedman test and post-hoc Dunn’s test, fraction of correct trials: expert, 92.7 ± 1.5% vs. contralateral trimming, 84.4 ± 1.7%, *P*=0.33; or vs. +ipsilateral trimming, 53.0 ± 1.3%, \*\*\**P*=0.0009). **d**-Acute injection of muscimol in S1 of expert animals reduced performance (*N*=5 mice, Wilcoxon rank sum test, fraction of correct trials: expert, 79.4 ± 1.9%, after muscimol, 52.5 ± 6.0%, \*\**P*=0.0079). **e**-Thin volumetric imaging was performed in order to correct *post-hoc* for axial brain motion artefacts during the task. Below, field of view containing mRuby2/GCaMP6s-expressing neurons. Single neurons were selected as regions of interest (ROIs) based on mRuby2 expression (inset), and recorded during training. Scale bar: 100*μ*m. **f**-Examples of average calcium responses to the P120_Go_ stimulus (green) and P280_No-go_ stimulus (red) from neurons in (e). Discrimination index values explained in (g) are indicated below each traces (DI, permutation test, *P*<0.05 for all these examples; see Methods). **g**-Example of ROC analysis for a neuron during an expert session. Left, probability distributions of the calcium responses for the P120_Go_ and P280_No-go_ trials. Right, ROC curve from which the DI is calculated. The DI represents the likelihood that the neuronal calcium response predicted which texture was presented. The significance level was determined using a permutation test (see Methods). **h**-The fraction of selective neurons before and after learning (*N*=12 mice, 875 neurons, naïve, 14.0 ± 2.6% (mean ± s.e.m.); expert, 25.0 ± 3.8%; Wilcoxon rank sum test, \**P*=0.03). **i**-Distribution of DIs in expert mice (*N*=12 mice, in green, P120_Go_ selective neurons: 146/875; in red, P280_No-go_ selective neurons: 57/875). **j**-The fraction of P120_Go_ (brown) and P280_No-go_ (orange) selective neurons before and after learning (*N*=12 mice, 875 neurons, Wilcoxon rank sum tests, P120_Go_ selective neurons: naïve, 9.5 ± 2.6%, expert, 19.2 ± 4.2%, \**P*=0.02; P280_No-go_ selective neurons: naïve, 5.3 ± 1.1%, expert, 7.2 ± 1.2%, *P*=0.33; P120_Go_ vs. P280_No-go_ selective fractions: naïve, *P*=0.36, expert, \*\**P*=0.006). (B to D, I and J) Lines represent individual mice and bars and error bars represent averages and s.e.m.

In order to monitor the stimulus selectivity of S1 neurons during texture discrimination learning, we co-expressed the genetically encoded calcium sensor GCaMP6s and the cell filler mRuby2, predominantly in excitatory L2/3 neurons using adeno-associated viral vectors (Fig. 1e, Supplementary Fig. 2)^20^. Single-cell responses were recorded using two-photon laser scanning microscopy (2PLSM) (Fig. 1e, f). Fast-volumetric imaging was used to allow for the correction of axial motion artifacts (Fig. 1e, see Methods)^21^.

Similar to previous studies^5, 14^, a fraction of the neurons displayed different response amplitudes upon presentation of the two textures (Fig. 1f and Supplementary Fig. 3). In order to determine the stimulus selectivity of individual neurons during learning we compared the calcium responses evoked by the two different textures and assessed their selectivity using a receiver-operating characteristic (ROC) curve analysis. This provided a discrimination index for each neuron (DI; Fig. 1g, see Methods)^22^. On average, the fraction of selective neurons increased with learning (Fig. 1h). Interestingly, we observed that in expert mice a larger fraction of the recorded population was selective for the P120 as compared to the P280 texture (Fig. 1i) and that this difference increased with learning (Fig. 1j).

The data indicate that the number of stimulus-selective L2/3 neurons increased with learning, with a bias toward one of the textures. There are three obvious explanations for this bias. First, it could have been caused by an increasing difference in the mouse’s behavior in response to both textures, which increased with learning (Supplementary Fig. 1). Second, L2/3 neurons could have an innate preference for coarser textures. Third, with learning, L2/3 neurons encode of higher-order features that are related to the textures (such as the stimulus-associated reward value or behavioral choice). To explore these possibilities, we conducted several experiments that allowed us to categorized neurons based on their activity in relation to the behavioral parameters, and we reassessed their selectivity after inverting the reward-contingencies.

### L2/3 neurons selectivity report sensory input and not behavioral output

We first investigated the possibility that the P120-selective neurons were primarily reporting a change in licking behavior that occurred with learning. As a first indication, we compared the delay between the calcium response to the texture presentation and to the 1^st^ lick response during hit trials. For the majority of neurons, the rise in the calcium trace occurred immediately after the texture presentation and preceded the 1^st^ lick with large jitter (Fig. 2a-c). This suggests that the activity of the P120-selective neurons was generated by the stimulus presentation itself but not by the licking response. However, this analysis could not exclude the possibility that selectivity had been influenced by an increasingly stereotyped pattern of behavioral responses during learning. To dissociate sensory from behavior-related neuronal activity we exposed the mice to the various task-related stimuli (i.e. sound cue, texture presentation and water reward) but presented them separately, without a structure, and prior to the training. We monitored the animal’s behavior (i.e. whisking and licking rates) (Fig. 2d). This pre-training session allowed us to categorize neurons based on their activity during task-related stimuli and behavior. We found that a large fraction of neurons (36.7% of the total population) exhibited touch-related activity during texture presentation while a few neurons were sensitive to auditory stimulation (0.8%) (Fig. 2e). Then we determined whether neurons reported whisking or licking-related signals. We trained a random forests machine learning model using the inferred firing rates from the calcium signal to assess for each neuron if its activity could predict whisking and/or licking rates. The model was trained using a range of positive and negative time lags of the neuronal activity relative to behavior, in order to account for possible pre-motor related activity (i.e. preceding the behavior) and/or sensory related activity (i.e. following the behavior). For each neuron we calculated the prediction power (PP), which reflected the correlation between the animals’ actual behavior and the behavior that was predicted by its activity (Fig. 2f). We plotted the PP distributions for whisking and licking rates as inferred from the GCaMP6s signal. This was compared to a control distribution that was inferred from the mRuby2 signal to assess the noise in PP measurement (Fig. 2g). We considered that neurons with a PP above the threshold criterion (5 × SD above the mean of the control distribution) were predictive of whisking and/or licking. We found that 9.4% of the neurons were partially predicting the animal’s whisking rate whereas only 2% predicted licking rates (Fig. 2f, h). We then compared the resulting categories with the selectivity that the neurons displayed in the texture discrimination task. We observed that most of the selective neurons found after training had been categorized as undefined or reporting touch (88%) under naïve conditions (Fig. 2i). Altogether these data strongly suggest that selective neurons did not signal whisking or licking behavior during the task. Moreover only 11% of the P120-selective neurons were predicting the animal’s whisking rate and 0% the licking rate. Thus, the biased increase in P120 selectivity during texture discrimination learning could not be explained by mere changes in the animal’s behavioral response.

**Fig. 2.**
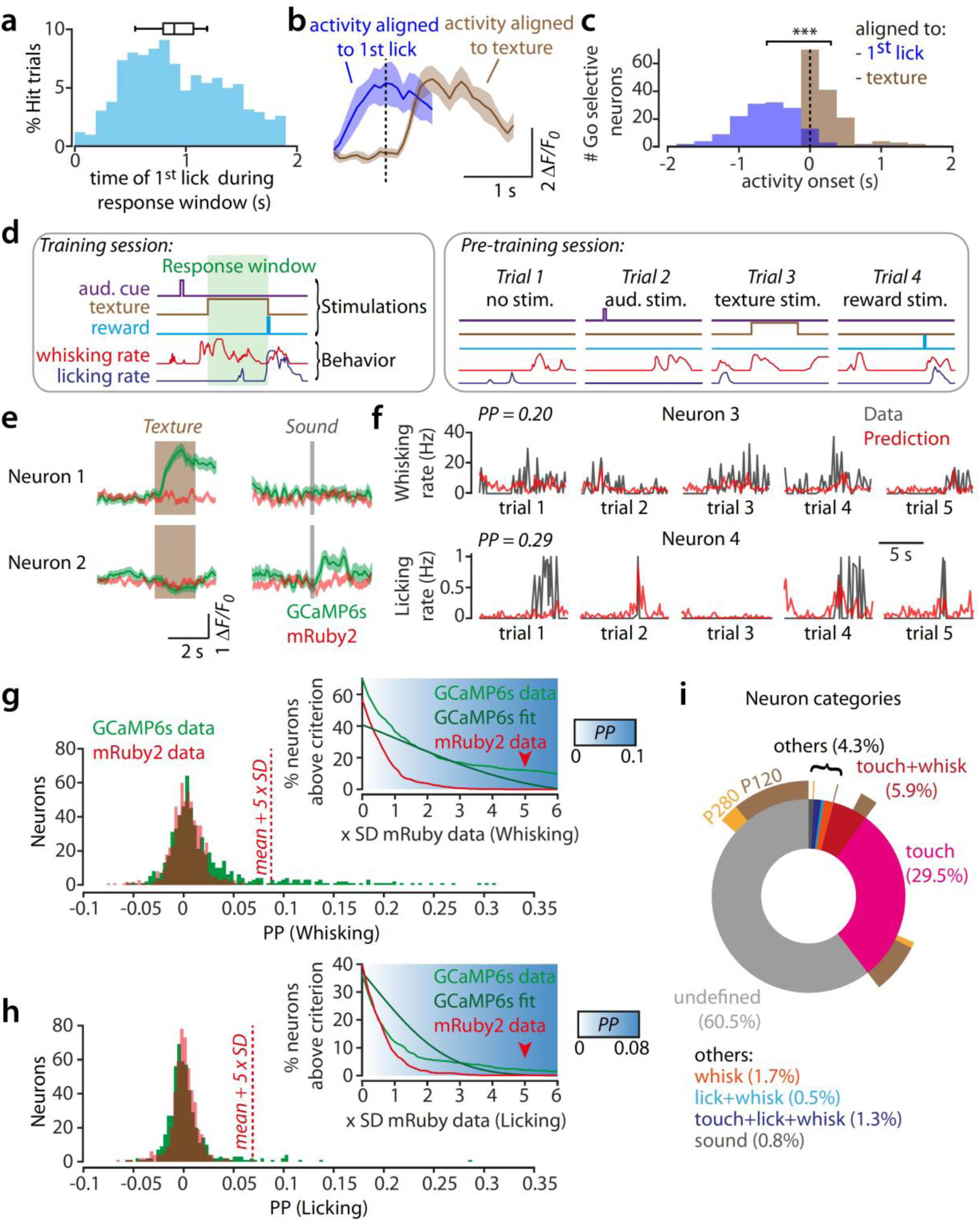
Activity of P120 selective neurons is not predicting licking behavior. **a**-Distribution of lick reaction times over all Hit trials across expert mice (*N*=923 trials). Box plot shows mean, 25^th^-75^th^ percentiles, and the minimum and maximum average reaction times over mice (*N*=12 mice). **b**-Example of the average calcium response in a P120 selective neuron of an expert mouse, aligned to texture presentation (brown) and the 1^st^ lick (blue). **c**-Distribution of the response onsets for all P120 selective neurons in expert sessions aligned to the texture presentation (brown) or to the 1^st^ lick (blue, *N*=12 mice, 146 neurons, average onset times: relative to texture, 0.29 ± 0.04 s; relative to 1^st^ lick, −0.61 ± 0.04 s, paired t-test, \*\*\**P*=2e-46). **d**-In a regular training session, stimuli and behavioral are tightly coupled, complicating the interpretation of the exact cause of the neuronal response. Therefore, prior to training, a random stimulus session was performed during which the 2-s texture presentation, the sound (auditory cue) or the water reward were delivered independently and in a pseudo-random fashion while recording spontaneous whisking and licking rates. Here shown, various trial examples in a random stimulus session as compared to a regular training session. **e**-Examples of two neurons significantly responding to texture or sound (green: GCaMP6s, red: mRuby2). Shaded areas represent s.e.m. **f**-Examples of two neurons decoding behavioral features as predicted by a Random-Forest decoder that was trained on inferred neuronal spike rates, and whisking and licking rates. Plots of whisking and licking rates (Data) superimposed with the scaled model prediction (Prediction). PP, prediction power. **g**-**h**-Prediction power (PP) distributions for whisking rates (G) and licking rates (H) reflecting the similarity between the behavioral data and the Random Forest decoding model based on the activity of individual neurons (inferred spikes; see Methods). The signal intensity fluctuation recorded from the mRuby2 channel was used as control. A neuron with a PP above 5 standard deviations (SD) of the control distribution was considered significantly predictive of the behavioral feature (*N*=591 neurons, 55 neurons predicted whisking rate and 10 neurons predicted licking behavior). The inset shows the percentage of neurons above a variable threshold criterion from the measured GCaMP6s PP values (light green) compared to normally distributed GCaMP6s PP values (dark green) and the measured mRuby2 PP values (red). This analysis shows that the fraction of neurons detected as predictive for whisking and licking is higher than a fraction obtained by chance for criterion values above 5 SD of the mRuby2 distribution. The variation of predictive fractions for whisking and licking around the chosen criterion of 5 SD (from 4 to 6 SD) of the mRuby2 distribution (indicated by arrows) is negligible. **i**-Percentage of neuron categories determined by combining outcomes of the response analysis in (e) and the behavioral decoding analysis in (f) (*N*=505 neurons, 8 mice). The outer layer of the pie chart represents the fraction of P120_Go_ and P280_No-go_ selective neurons within each of the categories.

### Stimulus selectivity is dynamic upon reversal learning

Next, we addressed whether the increase in the fraction of P120-selective neurons was based on an intrinsic bias to coarser textures, and/or whether it underlay the encoding of higher-order features that are related to the textures. These are conceivable possibilities. First, even though a preference is probably absent at the population level for the textures that we used, individual L2/3 neurons can display distinct preferences for certain textures^4^. Second, studies using comparable paradigms have reported that S1 neurons exhibit selectivity not only for the tactile stimulus but also for the behavioral choice^5, 14, 15^. However, such selectivity could result from the persistent coupling of stimulus and choice rather than the stimulus per se.

In order to test the two aforementioned possibilities, we uncoupled the behavioral outcome from the sensory stimulus by inverting the reward contingencies. To this end, expert mice were continued to be trained on the same textures where the P280 texture was rewarded and the P120 was not. Upon reversal the performance initially dropped to chance level (post-reversal naïve; Fig. 3a) before it reached the expert criterion again (post-reversal expert). We monitored neuronal activity throughout these phases and calculated their DIs. On average, we observed a net decrease of the neuronal selectivity during the post-reversal naïve phase as compared to the pre-reversal expert phase. This was followed by a net increase in selectivity during the post-reversal expert phase (Fig. 3b). By comparing, for each neuron, the DIs for the expert sessions before and after reversal we could define a variety of neuronal classes. For example, some neurons remained selective for the P120 texture, whereas others changed their selectivity to the P280, implying that they remained selective to the rewarded (Go)-texture (Fig. 3c). Altogether, 45% of the population had changed selectivity, whereas 55% had not (Fig. 3d).

**Fig. 3.**
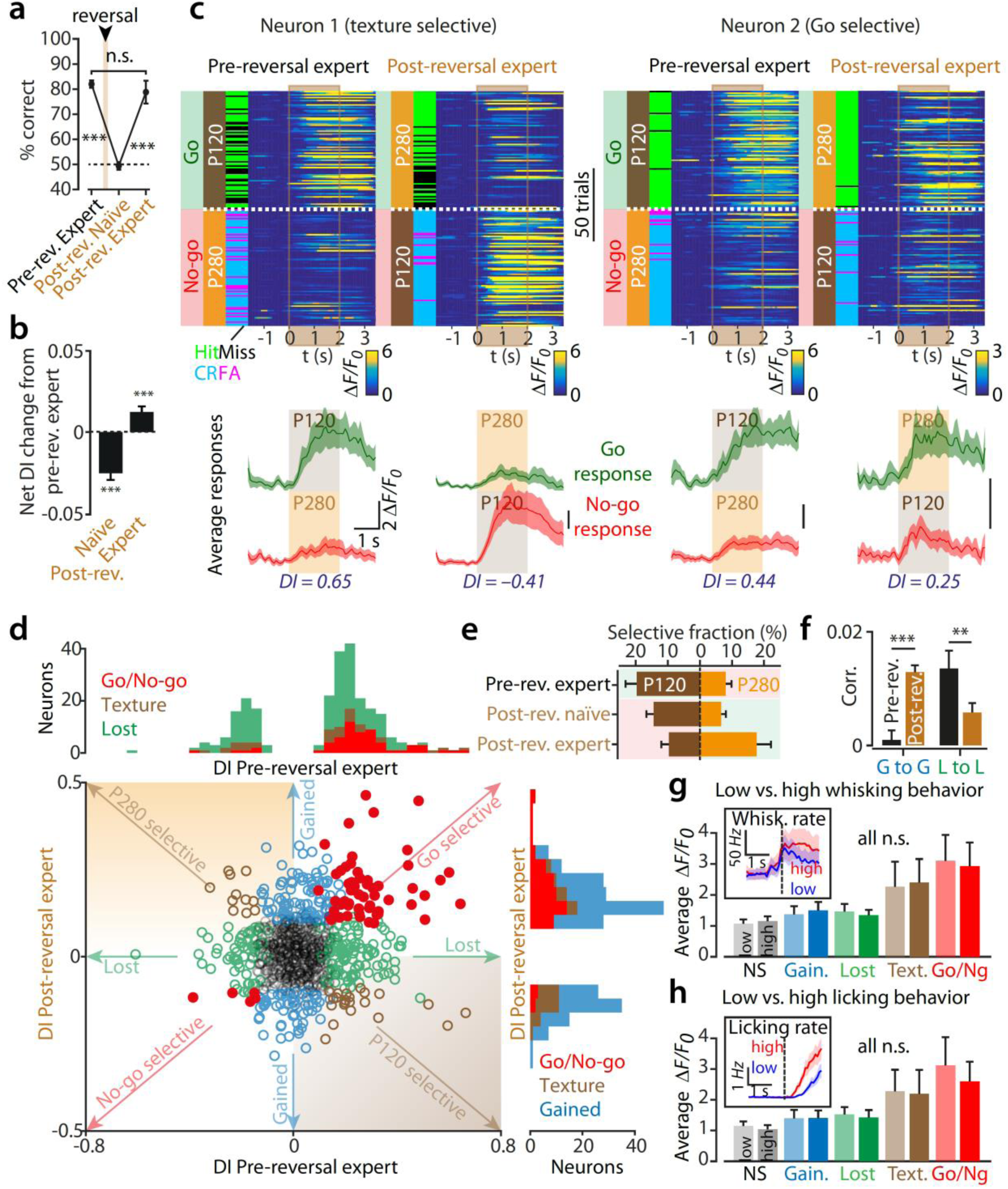
Origins of neuronal selectivity revealed by texture reversal learning. a-. Average performance of mice in pre-reversal expert, post-reversal naïve and post-reversal expert sessions (*N*=12 mice, rank sum test, pre-reversal expert vs. post-reversal naïve: \*\*\**P*=1.8e-4, post-reversal naïve vs. post-reversal expert: \*\*\**P*=3.3e-4, pre-reversal expert vs. post-reversal expert: *P*=0.62). **b**-Net DI change in post-reversal naïve and expert phases relative to their absolute pre-reversal expert DI. After reward inversion, selectivity of the population of neurons significantly reverts its sign, indicating that neurons preferentially respond to the previous Go texture P120 (*N*=863, t-test \*\*\**P*=9e-10). When mice become experts again after texture inversion, the selectivity of the neuronal population tends to revert back to the new Go stimulus (P280 texture; one sample t-test \*\*\**P*=6.6e-4). **c**-Single trial responses of two example neurons from expert sessions pre and post texture reversal aligned to texture presentation and sorted by stimulus type (160 trials from these expert sessions were displayed). Corresponding trial outcomes and performance are also shown. The neuron on the left remains selective for the P120 texture whereas the neuron on the right remains selective for the Go stimulus. Below, average calcium response with s.e.m. of these trials for P120 and P280 textures and their DI value indicated above. **d**-DIs of neurons in expert sessions pre vs. post reversal. Neurons in the top left and bottom right quadrants are texture selective (brown, *N*=33 neurons), as they preferentially respond to the same texture pre and post reversal. Neurons in the top right and bottom left quadrants are stimulus type selective (red, *N*=60 neurons), preferentially responding to either Go or No-go cue pre and post reversal. Other neurons lose (green, *N*=154 neurons) or gain (blue, *N*=144 neurons) selectivity, or remain non-selective (grey, *N*=412 neurons from 12 mice) throughout training. Top and right histograms show the distributions of DIs of selective neurons pre and post reversal. **e**-Average fractions of P120 and P280 selective neurons across mice before and after reversal learning (Pre-reversal expert: P120, 19.6 ± 3.6%, P280, 8.0 ± 1.8%; Post-reversal naïve: P120, 14.4 ± 2.3%, P280, 6.6 ± 1.4%; Post-reversal expert: P120, 9.6 ± 2.4%, P280, 17.6 ± 4.5%; *N*=12 mice, 863 neurons). **f**-Average spontaneous activity correlations between pairs of neurons that gained (G to G) or lost selectivity (L to L) comparing pre-reversal expert and post reversal expert (G to G: 224 pairs, \*\*\**P*=6e-9; L to L: 360 pairs, \*\**P*=0.0014 using paired t-tests). **g**-Effect of the magnitude of whisking during hit trials on calcium responses within each classes of neurons. Inset shows the average whisking rate across low (blue) and high (red) whisking rate trials after classification (see Methods). Graph shows the average calcium response during texture presentation across the various classes of neurons for low (light) and high (dark) whisking rates. Calcium responses were not influenced by whisking behavior (Paired t-test, P>0.69 for all classes, non-selective (NS): *N*=412 neurons, gained: *N*=144 neurons, lost: *N*=154 neurons, texture selective: *N*=33 neurons, Go/No-go selective (Go/Ng): *N*=60 neurons, across 12 mice). **h**-Same analysis as D with licking rate. Similarly, calcium responses were not influenced by licking behavior (Paired t-test, P>0.60 for all classes).

Since the population on average shifted its selectivity bias from P120 to P280 during the reversal (Fig. 3e), we can infer that it was not a priori preferring any of the two textures, which is in line with previous work^4^.

Within the dynamic population, a substantial fraction had lost or gained selectivity (19 and 18% respectively) upon reversal learning (Fig. 3d and Supplementary Fig. 4, which we classified as ‘lost’ and ‘gained’ neurons), indicating that sensory stimulus representation in the primary somatosensory cortex is largely dynamic. Such changes could be the result of network plasticity. To assess this, we calculated the level of co-fluctuation in spontaneous activity within each group, which may reflect the level of mutual connectivity^23, 24, 25^. Upon reversal, the level of co-fluctuation increased for gained neurons and decreased for lost neurons (Fig. 3f). This may indicate that reversal learning promotes the rewiring of local synaptic circuits.

We found two other classes of neurons, both of which remained selective after reversal learning (Fig. 3d). Some remained selective for the same texture (classified as texture neurons: 4.1%) while others changed their selectivity to the texture that represented the same reward contingency (classified as Go/No-go neurons: 7.5%).

There was no difference in the average calcium response for any of the classes above when comparing trials for which the animal displayed high whisking or licking rates with low rate trials (Fig. 3g, h, insets). This analysis suggests that the calcium response amplitude of the Go/No-go neurons, as well as for the other classes of neurons, did not depend on whisking (Fig. 3g) or licking activity (Fig. 3h). This result is in line with the decoding model (Fig. 2f-i) and indicates that the dynamics in calcium responses observed after reversal cannot be attributed to alterations in the behavioral response.

Altogether, the reversal learning experiment showed that the stimulus selectivity of L2/3 neurons in S1 was largely dynamic. The initial selectivity bias of the population towards the P120 texture was not caused by a higher sensitivity for coarser textures since it shifted towards the P280 texture after reversal. Instead, the selectivity of some neurons followed the reward contingency, suggesting that their activity is shaped by high-dimensional stimulus features.

### Go/No-go-selective neurons either signal upcoming behavioral choice or report stimulus-associated reward value

What determines the feature selectivity dynamics that we observed in the Go/No-go-selective class of neurons? We envision two possibilities. Neurons could report the upcoming choice, as previously described^5, 14, 15^. Alternatively, the reversal learning could have provoked a reassignment of neurons, shifting their selectivity from the formerly rewarded to the newly rewarded texture (from P120 to P280). The latter neurons would therefore signal the stimulus value rather than upcoming behavioral choice, as seen in other brain areas^8, 9, 26, 27^. To address this, we tracked the responses of individual Go/No-go-selective neurons according to the trial outcome (hits, misses, FAs and CRs) throughout the reversal learning process, distinguishing three phases: pre-reversal expert, post-reversal naïve, and post-reversal expert. Upon the reward contingency reversal, some neurons remained to have larger response during hit and FA trials as compared to misses and CR trials (e.g. Neuron 1 in Fig. 4a). Other cells exhibited larger responses during hits and misses for the original task, then showed larger responses during FA and CR trials in the naïve post-reversal phase, and finally regained response strength during hit and miss trials in the expert post-reversal phase (e.g. Neuron 2 in Fig. 4a). Thus, whereas the former neuron seemed to stably prefer a texture based on the final action-selection throughout all phases, the latter neuron seemed to update its selectivity during re-learning, possibly based on reward-value that was associated with the texture. We classified these two types of neurons by focusing on the post-reversal naïve phase in which the animals typically abandoned their previous behavioral strategy and made inconsistent choices. For each neuron we calculated a choice index (CI). Similar to the DI, this was based on a ROC curve analysis, but now comparing the response amplitudes between lick and no-lick trials (Fig. 4b). This analysis confirmed the existence of the two classes of Go/No-go-selective neurons (Fig. 4c), one for which the CI remained stable throughout the naïve phase after reversal, and one for which the CI was altered. Thus, while the former class was indicative of behavioral choice irrespective of the presented texture or licking behavior (classified as choice neurons; Fig. 4c, Supplementary Fig. 4), the latter was transiently uncoupled from choice and thus reported the value that was associated with the texture (i.e. indicative of the expected absence or presence of the upcoming reward; classified as value neurons; Fig. 4c). For both classes, the whisking and licking rates did not correlate with neuronal activity (Fig. 4d). In addition, only a few neurons in both classes were previously categorized as being predictive for whisking + licking, similar to the other classes of neurons (Fig. 4e). This confirms that the selectivity dynamics (or lack thereof) in choice and value neurons were not directly reporting alterations in behavior.

**Fig. 4.**
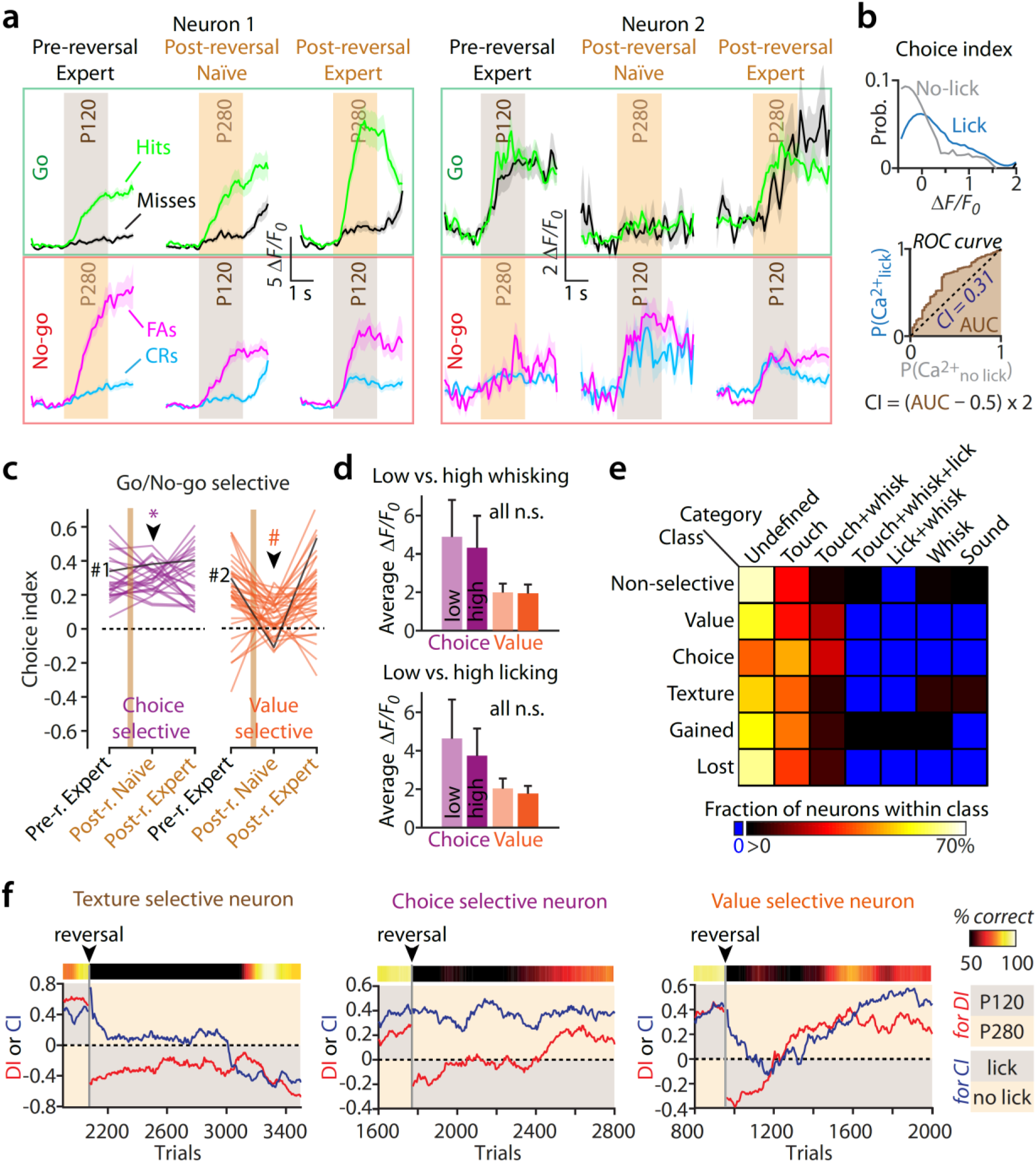
Go/No-go selective neurons signal choice or report stimulus-value. **a**-Average calcium responses in hits, misses, CRs and FAs of two Go selective neurons across reversal learning. For neuron 1, responses during hits and FAs remained high compared to misses and CRs reflecting the animal’s choice. For neuron 2, responses during hits and misses were higher before reversal and FAs and CRs were higher during the naïve phase of the reversal initially reflecting a selectivity for the P120 texture. But in the expert phase of reversal learning, responses were higher again during hits and misses reflecting a change in selectivity from the P120 to the P280 therefore signaling the value of the stimulus. **b**-Example of ROC analysis for calculating the choice index (CI). The calculation is similar to the DI except that calcium responses were sorted by the animal’s licking behavior (lick vs. no-lick trials). Top, probability distributions of the calcium responses for the lick and no-lick trials for an example neuron. Bottom, ROC curve from which the CI is calculated. The CI represent the likelihood that neuronal calcium response predicted whether the animal licked or not during the trial. The significance level was determined using a permutation test (see Methods). **c**-Changes in choice index (CI) of Go/No-go stimulus selective neurons in pre-reversal expert sessions, post-reversal naïve and expert sessions. Individual neurons CI were tested for significance, using random permutations, in the post-reversal naïve sessions. Neurons for which the CI was significant in the post-reversal naïve sessions and remained of the same sign as for pre- and post-reversal expert sessions (*) were classified as choice-selective neurons (purple). Other neurons for which the CI was either not significant or of the opposite sign in the post-reversal naïve sessions (#), as compared to pre- and post-reversal expert sessions, was classified as value-selective neurons (orange). **d**-Effect of the magnitude of whisking and licking during hit trials on calcium responses within choice and value neurons. Average calcium response during texture presentation across these two classes of neurons for low (light) and high (dark) whisking (top) and licking (bottom) rates. Calcium responses were neither influenced by whisking behavior (Paired t-test, P>0.83 for both classes, choice-selective: *N*=25 neurons, value-selective: *N*=35 neurons) nor licking behavior (Paired t-test, P>0.70 for both classes). **e**-Table showing the percentage of neurons of each categories within the different functional classes of neurons. Notably, none of the choice and value neurons was categorized as predictive for licking (number of neurons per classes for which corresponding categories were defined prior training: non-selective 271, value neurons 11, choice 14, texture 17, gained 90, lost 75). 0% is represented in blue in the look up table. **f**-Examples of temporal evolution of selectivity for textures (DI, red curve) and selectivity for licking behavior (CI, blue curve) across reversal learning for a texture-selective, a choice-selective and a value-selective neuron. For the texture-selective neuron, the DI is stably reporting the preference for the P120 texture while the CI is unstable. For the choice neuron, the CI signals the choice of the upcoming lick throughout training even in the naïve phase while the DI is unstable. For the value neuron, the CI is unstable throughout relearning while the DI is initially stably selective for P120 and then shifts its selectivity towards the P280 texture.

Altogether, these results show that Go/No-go selective neurons could be sub-divided into two classes: neurons that signaled the animal’s upcoming choice and neurons that reported the contextual stimulus value. To illustrate the differences between these classes, we provide examples of a texture, choice and value-selective neuron, showing the temporal evolution of their DI and CI throughout reversal learning (Fig. 4f). The DI of the texture-selective neuron reported a stable preference for the P120, while the CI changes signs. The CI of the choice-selective neurons reported a stable preference for the behavioral choice, while the DI changed signs. In contrast, both the DI and CI of the value-selective neuron were dynamic. Its CI gradually decreased to zero during the post-reversal naïve phase, while the DI immediately inversed sign and then slowly reverted back to the original sign. This confirms that the value-selective neuron initially remained to report the previously rewarded texture and then gradually reacquired a preference for the newly rewarded texture.

### Value-selective neurons display error history activity that precedes learning

Based on the preceding observations, we hypothesized that the gradual reacquisition of texture preference by the value-selective neurons carries a signal that predicts the upcoming improvement in the animal’s texture discrimination performance. Such a signal might consist of distinct response amplitudes during certain trials, which could depend on whether the animal had previously made correct or incorrect choices (Helmchen F. and Banerjee A., personal communication)^26, 28, 29^. Previous work suggests that a correct trial that follows an incorrect trial is considered more instructive for the animal than two consecutive correct trials^8, 9, 26^. To test this, we focused our analysis on those consecutive trials in which mice were actively licking upon texture presentation (i.e. hits and FAs), hence ensuring that they were engaged in the task. We compared the mean response amplitudes of hit trials that were preceded by a FA trial (*R_hit_*_(*post hit*)_) to those that were preceded by a hit trial (*R_hit_*_(*post hit*)_) (Fig. 5a). Trial sequences across mice were aligned to the point at which the reversal learning had reached the expert criterion (Fig. 5b and Supplementary Fig. 5). Averaged hit and FA rates separated from one another ∼140 trials before the expert criterion, indicating the moment at which mice started to improve their performance, termed learning onset (Fig. 5c, black arrow head). For non-selective, choice, texture, gained and lost selectivity neurons, we did not observe any difference between the *R_hit_*_(*post hit*)_ amplitudes and the *R_hit_*_(*post FA*)_ amplitudes (Fig. 5d). In contrast, for the contextual value-selective neurons, the average *R_hit_*_(*post FA*)_ response amplitudes became larger than the *R_hit_*_(*post hit*)_ amplitudes, at ∼260 trials before the expert criterion, and ∼120 trials before learning onset (Fig. 5c, d, red arrowhead). The two types of responses became similar again when mice performed above the expert criterion. During this interval, we did not observe a change in the sampling strategy of the texture confirming that the difference in responses is not associated with changes in licking and/or whisking rates (Supplementary Fig. 6). We used the normalized difference between *R_hit_*_(*post FA*)_ and *R_hit_*_(*post hit*)_ responses as an error history index (Fig. 5e), and observed that a large fraction of the value-selective neurons exhibited a transient increase in the error history as compared to the other neuronal classes that we had identified. Such an anticipation of the learning onset could not be deduced from the DI evolution (Supplementary Fig. 7). Altogether, these results indicate that the change in texture preference of value-selective neurons caries a signal that is indicative of the upcoming improvement in discrimination (i.e. learning).

**Fig. 5.**
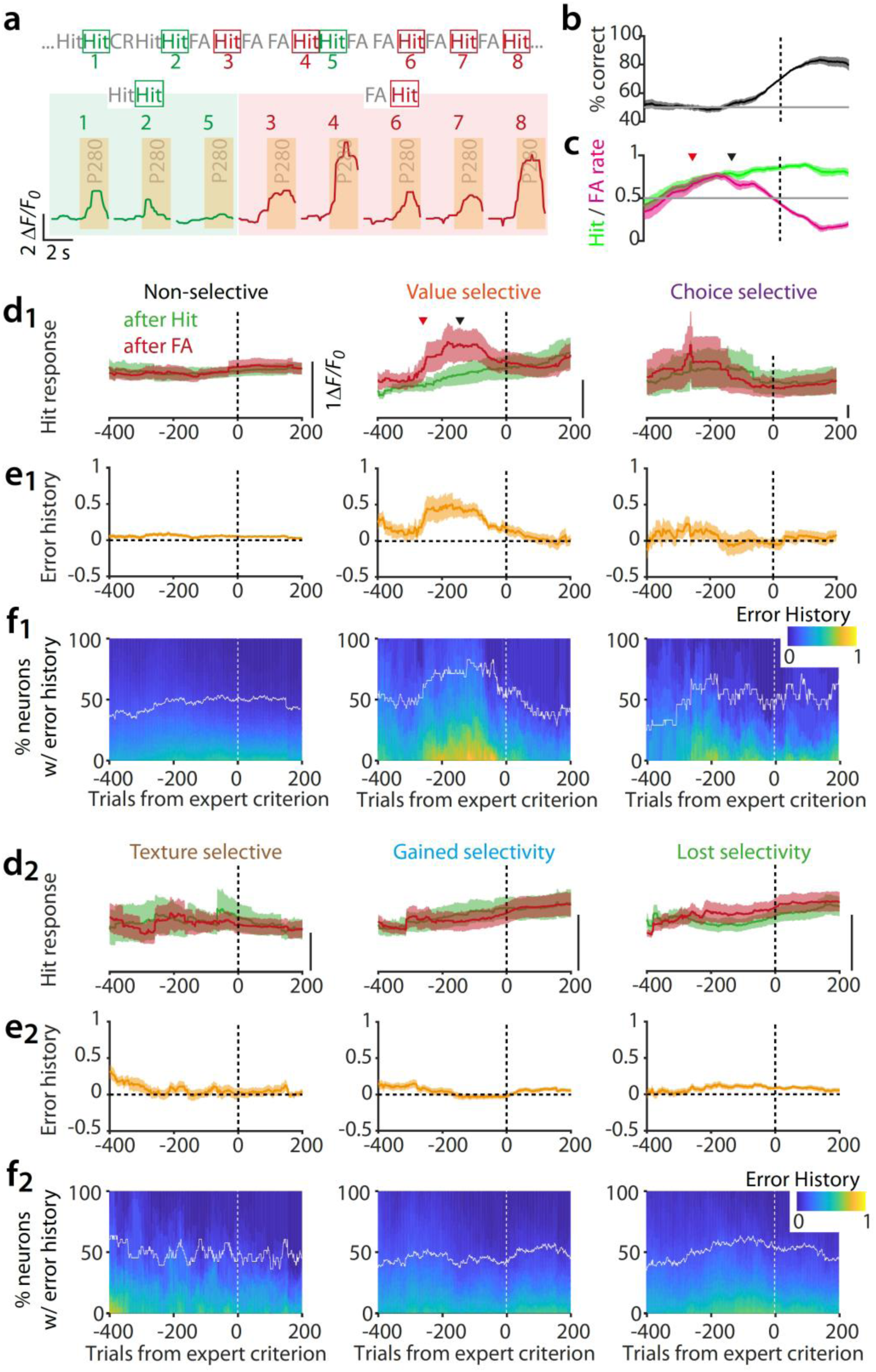
Behavioral error history is signaled by value-selective neurons during reversal learning prior to performance increase. **a**-Example of a value-selective neuron responses in Hit trials before the post-reversal performance increase. Hit trials preceded by a False Alarm trial (FA, dark red) contained larger responses than Hit trials that were preceded by another Hit trial (dark green). Numbers show the position of the Hit trials in the behavioral outcome sequence above. **b**-Average performance curves across mice realigned to the expert criterion (70% correct, dotted line) post-reversal, calculated with a rolling window of 200 trials (*N*=10 mice that displayed value-selective neurons). **c**-The corresponding average Hit and FA rate averages. (B to C) Shaded areas represent the 95% confidence intervals. **d**-The corresponding average calcium response (ΔF/F) in Hit trials that followed upon another Hit trial (green) or upon a FA trial (red) for non-selective (*N*=412 neurons from 12 mice), value-selective (*N*=35 neurons from 10 mice), choice-selective (*N*=24 neurons from 6 mice, **d_1_**), texture selective (*N*=33 neurons from 7 mice), gained (*N*=144 neurons from 12 mice) and lost selectivity (*N*=152 neurons from 12 mice, **d_2_**) neuronal classes. (**c** and **d**) Arrowheads indicate the trials at which the 95% confidence intervals separates for hit and FA rates (black) and the Hit-Hit and FA-Hit calcium responses (red). **e**-Calculated mean error history (see Methods). **f**-Percentage of neurons per class with significant error history rate (white line) above the cumulative distributions of error history values. (**b** to **e**) Shaded areas represent 95% CI.

## DISCUSSION

Previous studies have indicated that reward-based perceptual learning increases the reliability and selectivity of neuronal responses in primary sensory areas. As a consequence, the population activity stabilizes, which may improve perception^13, 16, 17, 18, 30^. We extend on this work by showing that task-related stimulus feature selectivity of neurons in the primary somatosensory cortex is dynamic upon reversal learning of a simple texture discrimination task. We found that a small fraction of the neurons is stably selective for texture or upcoming behavioral choice, while a large fraction loses, gains, or first loses and then regains selectivity when the reward contingencies of the stimuli are reversed (Fig. 3). This is remarkable, since upon reversal learning the capacity of the cortex to discriminate lower-order sensory features does not need to be altered in order for the mouse to perform the task. Nonetheless, the finding is congruent with the idea that learning continuously optimizes sensory representations in cortex, and strongly depends on the context^2, 13, 14^. In our study, the altered reward contingency could represent such a contextual change, since the animal has to suddenly shift its attention to the opposite texture to obtain a reward. Our data imply that the simple reversal of reward contingencies disorganizes the existing map, which is then extensively reshaped with relearning. The reshaping of this map could be the result of plasticity mechanisms that also underlie the experience-dependent tuning of neuronal response properties in primary sensory cortex^20^. In this case, Hebbian plasticity may drive the phenomenon, with the result that similarly tuned neurons become more strongly interconnected^23, 24, 25^. This is supported by our finding that both, neurons that gain selectivity and those that lose selectivity show higher co-fluctuation in spontaneous activity during the time they are selective (Fig. 3).

Similar to previous studies we found that selectivity is shaped by behavioral choice^5, 11, 14, 15^. For example, the population of neurons that became selective for the rewarded texture was larger than for the non-rewarded texture (Fig. 3). The reversal learning paradigm allowed us to assess the stability of this coupling, e.g. whether the Go-selective neurons stably respond to the rewarded texture, even during the post-reversal naïve phase, or whether they lose selectivity shortly upon reversal and then re-built it with relearning (Fig. 4). We find that more than half of the Go-selective neurons belonged to the latter class. Thus, their sensory responses were transiently uncoupled from the animal’s choice, and primarily depended on whether the presented texture was associated with the upcoming reward (or not). In future experiments it will be interesting to test whether repeated reversal learning would continue to renew the selective population, or whether the population would revert back to the original activity configuration.

Previous studies indicate that the value of a sensory stimulus is encoded by higher-order areas such as the posterior parietal, orbitofrontal, and retrosplenial cortices^8, 9, 26, 27^. Our data shows that value-encoding is also an attribute of a population of neurons in S1. The instructive cues for this selectivity could be manifold. For example, they could be provided by direct feedback from the aforementioned higher-order cortical areas, or they could be derived from sub-cortical areas that are implicated in attention and behavioral updating during learning^31^. Modulatory reinforcement signals that are associated with behavioral outcome could also play a major role^32, 33^. Indeed, reward-related response modulation has been observed in S1^28^, and was found to promote cortical plasticity processes related to visual response tuning in primary visual cortex^16^. We found that the value-selective neurons gradually regained their preference for the rewarded texture with relearning, which would be congruent with the idea that reward-related plasticity mechanisms contribute to shaping perceptual representations in cortex.

We also found that value neurons transiently displayed enhanced response amplitudes dependent on the animal’s behavioral error history (Fig. 5). During the naïve reversal phase these neurons showed higher responses in hit trials if the hit trial was preceded by a false alarm trial. This phenomenon was prominent during the transition from the naïve to expert reversal phase and forecasted the increase in behavioral performance. We speculate that the omission of reward-associated signals during a false alarm trial directs the animal’s attention towards the newly rewarded texture. Elevated attentional signals have been shown to modulate sensory-driven responses in visual cortex^34^. Thus, the attentional signals may be read out by the value neurons, which in turn reshape the texture selectivity of surrounding neurons. Together, this may enhance sensory perception.

## ACKNOWLEDGEMENTS

We thank Ariel Gilad for advice on the behavioral paradigm, Fritjof Helmchen and Abhishek Banerjee for discussions and suggesting the error history analysis, Jose Manuel Nunes for advice on assessment of selection criteria in the prediction model, Sebastien Pellat for technical support and engineering, Laura Bussien and Elodie Husi for assistance with histology. We thank Pieter Roelfsema for comments on the manuscript. This project was supported by the Swiss National Science Foundation (grants 31003A-153448, 31003A_173125, CRSII3_154453, and NCCR Synapsy 51NF40-158776), and a gift from a private foundation with public interest through the International Foundation for Research in Paraplegia.

## AUTHOR CONTRIBUTIONS

RC, SP and AH designed the experiments. RC and TK performed the experiments. RC analysed the data and LF performed random forests modelling. RC and AH wrote the manuscript. All of the authors edited the manuscript.

## MATERIALS AND METHODS

### Animals

All behavioral experiments were performed during the dark phase. All procedures were conducted in accordance with the guidelines of the Federal Food Safety and Veterinary Office of Switzerland and in agreement with the veterinary office of the Canton of Geneva (licence numbers GE/28/14, GE/61/17, and GE/74/18). C57BL/6J male mice (Janvier Labs) aged 6 weeks were group housed on a 12-hour light cycle (lights on at 8:00 AM) with littermates until surgery. Two weeks after surgery, animals were housed under standard conditions, with an inverted light-dark cycle 7-8 d before the first training session.

### Surgery and intrinsic optical imaging

Stereotaxic injections of adeno-associated viral (AAV) vectors were carried out on 6-week-old male C57BL/6 mice. A mix of O_2_ and 4% isoflurane at 0.4 l/min was used to induce anaesthesia followed by an intraperitoneal injection of MMF solution consisting of 0.2 mg/kg medetomidine (Dormitor, Orion Pharma), 5 mg/kg midazolam (Dormicum, Roche) and 0.05 mg/kg fentanyl (Fentanyl, Sinetica) diluted in sterile 0.9% NaCl. AAV1-hSyn1-mRuby2-GSG-P2A-GCaMP6s (Penn Vector Core; 100 nl)^20^ was delivered to L2/3 of the right C2 barrel-related column in S1 (1.4 mm posterior, 3.5 mm lateral from bregma, 300 µm below the pia). For long-term *in vivo* calcium imaging, a 3 mm diameter cranial window was implanted, as described previously^35^.

Two weeks after surgery, the C2 barrel column was mapped again using intrinsic optical imaging to confirm the location of mRuby2/GCaMP6s expression. To do this, a mix of O_2_ and 4% isoflurane at 0.4 l/min was used to induce anaesthesia followed by an intraperitoneal injection of MM solution consisting of 0.2 mg/kg medetomidine and 5 mg/kg midazolam diluted in sterile 0.9% NaCl. The C2 whisker was inserted into a capillary connected to a piezo actuator. Intrinsic signal from the C2 barrel column was collected during repeated whisker stimulation (1 s at 8 Hz). A 100-W halogen light source connected to a light guide system with a 700-nm interference filter was used to illuminate the cortical surface through the cranial window. Reflectance images 300 µm below the surface were acquired using a 2.7× objective and the Imager 3001F (Optical Imaging, Mountainside, NJ) equipped with a 256 × 256 pixels array charge-coupled device (CCD) camera. The built-in Imager 3001F analysis program was used to visualise the responses and produce an intrinsic signal image by dividing the stimulus signal by the pre-stimulus baseline signal. An image of the vasculature was then acquired using a 546-nm interference filter, and superimposed to the intrinsic signal image. This reference image was used later to select an appropriate field of view (FOV) using 2PLSM. After this procedure, a metal post was implanted laterally to the window using dental acrylic to restrict head movement during behavior and imaging.

### Habituation and water deprivation

Mice were handled and accustomed to be head restrained on the training setup for 10-15 min during 4-5 d. Water deprivation started 3-5 d before the first training session and ceased at the end of the training. Weight was monitored daily during this period and the amount of water given was adjusted to prevent them from losing more than 15% of their original weight. Altogether, mice received a minimum of 1 ml of water per day corresponding to the amount they drank during the training as rewards plus the amount that the experimenter provided outside of the training sessions.

### Texture discrimination task

Mice were trained to discriminate between two commercial-grade sandpapers (P120 and P280) in a Go/No-go paradigm as described previously^5^. The control of the devices and the recording of behavioural parameters were performed using a data acquisition interface (PCI 6503, National Instruments) and custom-written LabWindows/CVI software (National Instruments). Licks were detected electrically. Mice remained on a metallic plate that maintained an electrical potential difference with the licking spout. The electrical circuit was closed when mice touched the spout with their tongue, producing a 1.2-µA current that was detected by the acquisition interface. Whisking activity was measured with an optical barrier that detected the changes in intensity when whiskers swept through it at a sampling frequency of 10 kHz. To achieve this, an 850 nm LED beam was used as light source (HIR204C, Everlight Electronics) and an 860 nm phototransistor (PT 202C, Everlight Electronics) was used to detect intensity variations through a 1-mm hole placed 60 mm away from the light source. Whisking activity was quantified as the frequency at which individual whiskers crossed the light beam placed ∼1 mm in front of and centered on the presented texture. The licking and whisking rates were calculated as the average number of events over a sliding window of 100 ms and normalized per second.

Sandpapers were attached onto a four-arm wheel (2 × 2 of the same sandpapers) mounted on a stepper motor (T-NM17A04, Zaber) and a motorized linear stage (T-LSM100A, Zaber) to move textures in and out of reach of whiskers. At the start of each trial, the wheel spun for a random amount of time in the back position of the linear stage (approximately between 0.5 and 1 s) and stopped between 2 textures positions. To present a texture the linear stage first moved to the front position and then the stepper motor rapidly slid the sandpaper into the whisker field at approximately 15 mm from the snout with an angle of 70° relative to the rostro-caudal axis. In the first phase of the training, the coarser P120 sandpaper was the Go-stimulus (i.e. target stimulus for which the mouse was incited to lick the spout in order to receive a water reward) and the P280 sandpaper was the No-go stimulus (i.e. non-target stimulus for which the mouse was incited to refrain from licking the spout). Initially, mice were trained to trigger a 4-6 µl sucrose water reward (100 mg/ml) by licking the spout during the presentation of the Go-stimulus (P120). Then, they were gradually familiarized with the No-go stimulus (from 0 to 30% of the trials) within two sessions (one session per day, 150-300 trials per session). Imaging started when Go and No-go stimuli were pseudo-randomly presented with 50% probability for each trial type with a maximum of 4 consecutive presentations of the same stimuli. A trial consisted of a 1-s pre-stimulus period followed by a 3-kHz auditory cue for 200 ms, a delay period of 500 ms after which the texture reached the whiskers within 150 ms and remained there for 2 s before being retracted. Licking during the Go stimulus presentation triggered a water reward at the end of the 2-s of presentation, and the corresponding trial was scored as a ‘hit’. Licking during the No-go stimulus presentation triggered a 500-ms white noise at the end of the 2-s of presentation plus a 5-s time out period and the trial was scored as a ‘false alarm’ (FA). In the absence of a lick during stimulus presentation, trials were scored as a ‘miss’ or a ‘correct rejection’ (CR) for Go and No-go stimuli respectively. To prevent the mice from compulsive licking during training, in addition to the aforementioned rules, mice had to show a 2-fold increase in the licking rate during stimulus presentation as compared to the pre-stimulus baseline period to get rewarded on Go stimulus presentation. Around 250-400 trials per session were performed (1 session per day) at a rate of ∼6 trials/min.

The overall performance of the animal was calculated as the percentage of correct trials (hits + CRs) over an entire session or over a sliding window of 200 trials (Fig. 3 and Supplementary Fig. 5). The hit and FA rates were calculated as *N_hit_*/(*N_hit_* + *N_miss_*) and *N_FA_* /(*N_FA_* + *N_CR_*) respectively where *N* is a number of trials for an entire session or over a sliding window of 200 trials (Fig. 3 and Supplementary Fig. 5). Mice were considered experts when the average performance per session reached a level of 70% correct trials (the expert criterion) over two consecutive sessions. In the second phase of training (i.e. reversal learning), Go and No-go stimuli were inverted (P280 and P120 respectively) and mice were trained until they reached the same expert criterion again in two consecutive sessions.

### 2PLSM

We used a custom built 2-photon laser scanning microscope mounted onto a modular *in vivo* multiphoton microscopy system (https://www.janelia.org/open-science/mimms-10-2016) equipped with an 8-kHz resonant scanner and a 16× 0.8NA objective (Nikon, CFI75), and controlled with Scanimage 2016b^36^ (http://www.scanimage.org). Fluorophores were excited using a Ti:Sapphire laser (Chameleon Ultra, Coherent) tuned to *λ* = 980 nm that was slightly underfilling the back aperture of the objective to extend the depth of field to 5 µm. Fluorescent signals were collected with GaAsP photomultiplier tubes (10770PB-40, Hamamatsu) separating mRuby2 and GCaMP6s signals with a dichroic mirror (565dcxr, Chroma) and emission filters (ET620/60m and ET525/50m, respectively, Chroma). Fast volumetric imaging was performed at 11.5 Hz using a piezo z-scanner (P-725 PIFOC, Physik Instrumente) for moving the objective over the z-axis. Each acquisition volume consisted of 5 contiguous planes (with 5-µm steps between planes) of 400 × 400 µm (512 × 256 pixels) allowing post-hoc z-motion correction which may be generated by licking-induced brain motion artifacts^21^.

### Image processing

Image acquisitions were processed using custom-written MATLAB scripts and ImageJ (http://rsbweb.nih.gov/ij/). Lateral and axial motion corrections were performed using the mRuby2 signal as a reference. First, rigid lateral movement vectors were calculated based on individual trial movies from the average z-projection of the 20-µm imaged volumes using the NoRMCorre MATLAB toolbox^37^. Residual bidirectional scanning artifact vectors were calculated using a highest-pixel-line signal correlation between the two scanning directions on the entire frame. Inter-trial registration was calculated using a custom-written cross-correlation algorithm based on the rigid image stack registration plugin in ImageJ. All calculated lateral motion corrections were applied on both the mRuby2 and GCaMP6s signals. Second, axial motion correction was performed using cross-correlation on linearly interpolated volumes (with a factor 3). The image planes with the highest correlation to a reference image, defined as the center image plane of the first volume, were selected. For an unbiased extraction of the GCaMP6s fluorescence signals from individual neurons, regions of interest (ROIs) were drawn manually for each session based on neuronal shape using the mRuby2 signal. The fluorescence time-course of each neuron (*F_measured_*) was measured as the average of all pixel values of the GCaMP6s signal within the ROI. Local neuropil signal (*F_neuropil_*) was measured for each ROI as the average of pixel values within an automatically defined ring of 15 µm width, 2 µm away from the ROI (excluding overlap with surrounding ROIs)^38^. The fluorescence signal of a cell body was then estimated as *F*(*t*) = *F_measured_* (*t*) − *r* × *F_neuropil_* (*t*) with *r* = 0.7 ^39^. Residual trends were removed by subtracting the 8^th^ percentile of each trial ^40^. Normalized calcium traces Δ*F*/*F*_0_ were calculated as (*F* − *F*_0_)/*F*_0_, where *F*_0_ is the median of the individual mean baseline fluorescence signal of each trial over a 1-s period before the start of the stimulation. For individual stimulation sessions (see *Individual stimulation session and neuron categorization* section) and spontaneous activity recordings, *F*_0_ is the 30^th^ percentile of each trial trace. For display, traces were additionally filtered with a Savitzky-Golay function (2^nd^ order, 500-ms span).

### Activity onset analysis

Normalized calcium traces (Δ*F*/*F*_0_) were aligned to either the onset of the texture presentation or to the first lick during the texture presentation for each neuron across all hit trials of an expert session. For both realignments, the onset of the neuronal response was calculated as the time, relative to the texture or first lick onset, at which the average of the response reached half of its maximum amplitude.

### Individual stimulation session and neuron categorisation

Prior to the start of the training, 9 mice were imaged in the experimental training configuration, where task stimulation features were presented independently in a pseudo-random fashion. Data acquisition was organized in trials of 10 s, each starting with a 3-s baseline after which one of the following conditions was presented at a random time within a 4-s window: 2-s texture, 0.2-s sound (auditory cue) or water valve opening to incite licking, and finishing with another 3-s of recording. In 20% of the trials, no stimulation was applied. Whisking and licking events were recorded during the course of the session.

To determine if neuronal activity was significantly modulated by texture or sound stimulations, we compared, for each neuron across trials, the average normalized fluorescence during 1 s before and after stimulus onset using a paired-sample t-test at a significance threshold of 5%. To account for noise in our data due to possible stimulation-induced movement artefacts, we performed the same test using the mRuby2 signal. None of the neuron showed a significant change in mRuby2 signal upon texture and sound stimulations.

We used a random forests machine-learning algorithm to decode behavioural features (licking and whisking rates) from the activity of single neurons. This procedure allowed us to categorize single neurons as either decoding whisking, licking or both. Given the slow kinetics of calcium transients captured by the GCaMP6s sensor, spiking rates were inferred from the Δ*F*/*F*_0_ trace and used as input to the algorithm, which allowed to temporally match behavioral event variations (i.e. whisking or licking rates) to neuronal activity. Firing rates at each imaging frame were inferred from normalized calcium traces (Δ*F*/*F*_0_) using a fast nonnegative deconvolution method (https://github.com/jovo/oopsi)^41^ with variable background fluorescence estimation and a *K_d_* of 144 nM^42^. In order for the algorithm to capture differences in activity levels between neurons, all trial traces of all neurons recorded per mouse were concatenated before inferring spikes. To account for putatively preceding pre-motor and/or following sensory activity relative to behavioural events in S1, neuronal activity was shifted negatively and positively in time with a maximum shift of 500 ms. Eleven time bins of inferred firing rates (discretized in time bins of 100 ms) centred on zero time-shift were used to predict instantaneous behavioural features and composed a vector *X_i_*(*t*) = [*x_i_*(*t*) − 500 *ms*), … *x_i_*(*t*), …, *x_i_*(*t* + 500 *ms*)] where *x_i_*(*t*) represents the inferred firing rates of the *i*th neuron at zero time-shift. Licking and whisking rates were down sampled to 11.5 Hz in order to temporally match calcium imaging data. The *ranger* function of the ranger R package version 0.10.1 was used to construct regression forests, with each behavioural feature as dependent variable and the binned inferred firing rates of a given neuron as predictors. For each neuron, two regression forests were constructed, one to decode whisking and the other licking. Most arguments of the function were kept at default settings, except the following: the number of trees was set to 128, the minimum size of terminal nodes was set to 2, the number of predictor variables randomly sampled at each node split was set to the maximum between 1 or the third of the number of predictors, and the variable importance mode was set to “impurity”. To obtain a prediction for all trials, 5-fold cross-validation was applied by training the algorithm on 80% of the trials (i.e. training set) and evaluating it on the remaining 20% of the trials (i.e. test set). Since data acquisition was discretized by trial, for each cross-validation the training and test set trials were concatenated for training and prediction, respectively. For each neuron and for each behavioural feature, the decoding accuracy was assessed by computing the Pearson’s product-moment correlation coefficient between the observed and predicted behavioural event fluctuations. In order to get an estimate of the noise in the prediction levels, the same analysis was performed using the mRuby2 signal as a control. Neurons were classified as decoding a given behavioural feature if their Pearson’s correlation coefficient computed on the GCaMP6s signal was five standard deviations away from the mean of the Pearson’s correlation coefficients for all neurons computed on the mRuby2 signal. Neurons meeting these criteria for both whisking and licking were classified as decoding both behavioural features.

### Spontaneous activity correlation

Spontaneous calcium transients were recorded for 10 min after mice reached the expert level before and after texture reversal. Pairwise Pearson’s correlation coefficients were calculated on the normalized calcium traces.

### Discrimination and choice indices

The selectivity of each neuron was expressed by a Discrimination index (DI) that was calculated based on neurometric functions using a receiver operating characteristic (ROC) analysis^22, 43, 44^. Normalized mean calcium responses (Δ*F*/*F*_0_) during the 2 s stimulus presentations in the P120 texture trials were compared to the P280 texture trials. ROC curves were generated by plotting, for all threshold levels, the fraction of P120 trials against the fraction of P280 trials for which the response exceeded threshold. Threshold levels were defined as a linear function from the minimal to the maximal calcium responses. DI was computed from the area under the ROC curve (AUC) as follows: *DI* = (*AUC*− 0.5) × 2. DI values vary between −1 and 1. Positive or negative values indicate larger or smaller responses to P120 than to P280 texture presentations, respectively. Statistical significance of the measured DI value was assessed by performing a permutation test, from which a sampling distribution was obtained by shuffling the texture labels of the trials 10,000 times. The measured DI was considered significant when it was outside of the 2.5^th^ – 97.5^th^ percentiles interval of the sampling distribution. For the choice index (CI), the same calculation was performed, with the difference that trials where the animal licked during the response window were compared to trials with no lick. For building the temporal evolution of the DI and CI across reversal learning, both indices were calculated on the sliding window of 100 trials every 5 trials.

### Calcium responses relative to behavioral strategies

For all hit trials of an expert session, average whisking and licking rates were calculated as the average number of events over the entire response window. For each mouse, the median value in both distributions was used to separate low and high whisking or licking rate trials.

### Error history

Error history for each neuron was calculated as the normalized difference between the average calcium response during ‘hits’ (*R̅*) over a sliding window of 200 trials as follows:

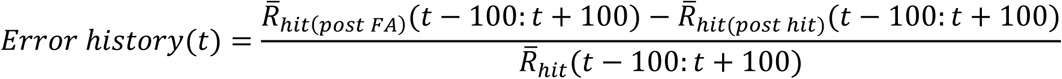

where *R_hit_*_(*post FA*)_ is a ‘hit’ response preceded by a ‘FA’, *R_hit_*_(*post hit*)_ is a ‘hit’ response preceded by another ‘hit’, *R_hit_* is any ‘hit’ response and *t* is the trial position along the learning sequence. To estimate the fraction of neurons with an error history above chance over reversal learning, for each neuron, all ‘hit’ responses within each window of 200 trials were randomly permuted, replacing *R_hit_*_(*post FA*)_, *R_hit_*_(*post hit*)_ and *R_hit_* in their respective trial positions and an error history value was calculated. This process was repeated 1000 times to obtain 95% confidence intervals for each observed error history value.

### Immunohistochemistry

Post-hoc immunohistochemistry of GABA was performed on mRuby2/GCaMP6s expressing neurons. 100-µm thick tangential sections were produced using a vibratome (Leica VT 1000). The sections were washed 3 × 3 minutes in 500 µl Tris-buffered saline (0.1 M Tris, 150 mM NaCl) containing 0.1% Tween (TBST), then pre-treated with TBST and 0.1% Triton-X for 20 minutes followed by a 3 × 3 minute TBST wash. They were blocked in 300 µl TBST containing 10% normal donkey serum (ab7475, Abcam) for 1 hour and incubated with mouse anti-GABA antibody (ab86186, Abcam) diluted 1:500 for 72 hours at 4°C. After another 5 × 3 minutes wash in TBST they were incubated in 300 µl of donkey anti-mouse antibody coupled to Alexa Fluor 647 (A32787, Thermo Fisher Scientific) diluted 1:200 in TBST for one hour at room temperature. Finally, they were washed 10 × 3 minutes in TBST and then for one hour in PBS before being mounted onto glass slides. We applied Fluoroshield mounting medium with DAPI (Abcam) before applying the coverslip. The sections were imaged using a Zeiss Confocal LSM800 Airyscan.

### S1 inactivation and whisker trimming

To inactivate S1, the GABA (G-aminobutyric acid) agonist muscimol was injected in expert mice. During the test session, high baseline performance (>70%) was first recorded for 100 trials before the injection was performed. Under light anesthesia (4% isoflurane at 0.4 l/min), a small hole was drilled through the imaging window above the previously mapped C2 barrel column to provide access to a glass pipette through which 300 nl of Muscimol (Bodipy-TMR-X, 5mM in cortex buffer with 5% DMSO, Thermo Fisher Scientific) was injected at 300 and 500 µm below the pia. Mice were left to recover for 45 min and their behavioural performance was then assessed for another 100 trials. For the whisker trimming experiment, a similar baseline performance was first recorded for 100 trials before trimming the whiskers on the side of the snout contralateral to the texture presentations, and tested the performance for 50 trials. This ensured that trimming itself did not alter performance. Then, the whiskers that were in contact with the textures (ipsilateral to the texture presentation side) were trimmed, and the effect on task performance was measured for another 50 trials.

### Statistics

All statistics were performed using MATLAB. For all figures, significance levels were denoted as \**P*<0.05, \*\**P*<0.01, \*\*\**P*<0.001. No statistical methods were used to estimate sample sizes. Non-parametric tests were used for sample sizes smaller than 15.

## REFERENCES

1. Holtmaat A, Caroni P. Functional and structural underpinnings of neuronal assembly formation in learning. Nat Neurosci 19, 1553–1562 (2016).

2. Makino H, Hwang EJ, Hedrick NG, Komiyama T. Circuit Mechanisms of Sensorimotor Learning. Neuron 92, 705–721 (2016).

3. Safaai H, von Heimendahl M, Sorando JM, Diamond ME, Maravall M. Coordinated population activity underlying texture discrimination in rat barrel cortex. J Neurosci 33, 5843–5855 (2013).

4. Garion L, et al. Texture coarseness responsive neurons and their mapping in layer 2-3 of the rat barrel cortex in vivo. Elife 3, e03405 (2014).

5. Chen JL, Carta S, Soldado-Magraner J, Schneider BL, Helmchen F. Behaviour-dependent recruitment of long-range projection neurons in somatosensory cortex. Nature 499, 336–340 (2013).

6. Hubel DH, Wiesel TN. Receptive fields of single neurones in the cat’s striate cortex. J Physiol 148, 574–591 (1959).

7. Stiebler I, Neulist R, Fichtel I, Ehret G. The auditory cortex of the house mouse: left-right differences, tonotopic organization and quantitative analysis of frequency representation. J Comp Physiol A 181, 559–571 (1997).

8. Akrami A, Kopec CD, Diamond ME, Brody CD. Posterior parietal cortex represents sensory history and mediates its effects on behaviour. Nature 554, 368–372 (2018).

9. Hwang EJ, Dahlen JE, Mukundan M, Komiyama T. History-based action selection bias in posterior parietal cortex. Nat Commun 8, 1242 (2017).

10. Felleman DJ, Van Essen DC. Distributed hierarchical processing in the primate cerebral cortex. Cereb Cortex 1, 1–47 (1991).

11. Pho GN, Goard MJ, Woodson J, Crawford B, Sur M. Task-dependent representations of stimulus and choice in mouse parietal cortex. Nat Commun 9, 2596 (2018).

12. Francis NA, Winkowski DE, Sheikhattar A, Armengol K, Babadi B, Kanold PO. Small Networks Encode Decision-Making in Primary Auditory Cortex. Neuron 97, 885–897 e886 (2018).

13. Poort J, et al. Learning Enhances Sensory and Multiple Non-sensory Representations in Primary Visual Cortex. Neuron 86, 1478–1490 (2015).

14. Chen JL, Margolis DJ, Stankov A, Sumanovski LT, Schneider BL, Helmchen F. Pathway-specific reorganization of projection neurons in somatosensory cortex during learning. Nat Neurosci 18, 1101–1108 (2015).

15. Yang H, Kwon SE, Severson KS, O’Connor DH. Origins of choice-related activity in mouse somatosensory cortex. Nat Neurosci 19, 127–134 (2016).

16. Goltstein PM, Coffey EB, Roelfsema PR, Pennartz CM. In vivo two-photon Ca2+ imaging reveals selective reward effects on stimulus-specific assemblies in mouse visual cortex. J Neurosci 33, 11540–11555 (2013).

17. Pecka M, Han Y, Sader E, Mrsic-Flogel TD. Experience-dependent specialization of receptive field surround for selective coding of natural scenes. Neuron 84, 457–469 (2014).

18. Schoups A, Vogels R, Qian N, Orban G. Practising orientation identification improves orientation coding in V1 neurons. Nature 412, 549–553 (2001).

19. Huber D, et al. Multiple dynamic representations in the motor cortex during sensorimotor learning. Nature 484, 473–478 (2012).

20. Rose T, Jaepel J, Hubener M, Bonhoeffer T. Cell-specific restoration of stimulus preference after monocular deprivation in the visual cortex. Science 352, 1319–1322 (2016).

21. Chen JL, Pfaffli OA, Voigt FF, Margolis DJ, Helmchen F. Online correction of licking-induced brain motion during two-photon imaging with a tunable lens. J Physiol 591, 4689–4698 (2013).

22. Takahashi N, Oertner TG, Hegemann P, Larkum ME. Active cortical dendrites modulate perception. Science 354, 1587–1590 (2016).

23. Ko H, Hofer SB, Pichler B, Buchanan KA, Sjostrom PJ, Mrsic-Flogel TD. Functional specificity of local synaptic connections in neocortical networks. Nature 473, 87–91 (2011).

24. Cossell L, et al. Functional organization of excitatory synaptic strength in primary visual cortex. Nature 518, 399–403 (2015).

25. Cohen MR, Kohn A. Measuring and interpreting neuronal correlations. Nat Neurosci 14, 811–819 (2011).

26. Hattori R, Danskin B, Babic Z, Mlynaryk N, Komiyama T. Area-Specificity and Plasticity of History-Dependent Value Coding During Learning. Cell, (2019).

27. Sul JH, Kim H, Huh N, Lee D, Jung MW. Distinct roles of rodent orbitofrontal and medial prefrontal cortex in decision making. Neuron 66, 449–460 (2010).

28. Lacefield CO, Pnevmatikakis EA, Paninski L, Bruno RM. Reinforcement Learning Recruits Somata and Apical Dendrites across Layers of Primary Sensory Cortex. Cell Rep 26, 2000–2008 e2002 (2019).

29. Campagner D, et al. Prediction of Choice from Competing Mechanosensory and Choice-Memory Cues during Active Tactile Decision Making. J Neurosci 39, 3921–3933 (2019).

30. Peron SP, Freeman J, Iyer V, Guo C, Svoboda K. A Cellular Resolution Map of Barrel Cortex Activity during Tactile Behavior. Neuron 86, 783–799 (2015).

31. Wimmer RD, Schmitt LI, Davidson TJ, Nakajima M, Deisseroth K, Halassa MM. Thalamic control of sensory selection in divided attention. Nature 526, 705–709 (2015).

32. Watabe-Uchida M, Eshel N, Uchida N. Neural Circuitry of Reward Prediction Error. Annu Rev Neurosci 40, 373–394 (2017).

33. Pi HJ, Hangya B, Kvitsiani D, Sanders JI, Huang ZJ, Kepecs A. Cortical interneurons that specialize in disinhibitory control. Nature 503, 521–524 (2013).

34. Pooresmaeili A, Poort J, Roelfsema PR. Simultaneous selection by object-based attention in visual and frontal cortex. Proc Natl Acad Sci U S A 111, 6467–6472 (2014).

35. Holtmaat A, et al. Long-term, high-resolution imaging in the mouse neocortex through a chronic cranial window. Nat Protoc 4, 1128–1144 (2009).

36. Pologruto TA, Sabatini BL, Svoboda K. ScanImage: flexible software for operating laser scanning microscopes. Biomed Eng Online 2, 13 (2003).

37. Pnevmatikakis EA, Giovannucci A. NoRMCorre: An online algorithm for piecewise rigid motion correction of calcium imaging data. J Neurosci Methods 291, 83–94 (2017).

38. Kerlin AM, Andermann ML, Berezovskii VK, Reid RC. Broadly tuned response properties of diverse inhibitory neuron subtypes in mouse visual cortex. Neuron 67, 858–871 (2010).

39. Chen TW, et al. Ultrasensitive fluorescent proteins for imaging neuronal activity. Nature 499, 295–300 (2013).

40. Dombeck DA, Khabbaz AN, Collman F, Adelman TL, Tank DW. Imaging large-scale neural activity with cellular resolution in awake, mobile mice. Neuron 56, 43–57 (2007).

41. Vogelstein JT, et al. Fast nonnegative deconvolution for spike train inference from population calcium imaging. J Neurophysiol 104, 3691–3704 (2010).

42. Lock JT, Parker I, Smith IF. A comparison of fluorescent Ca(2)(+) indicators for imaging local Ca(2)(+) signals in cultured cells. Cell Calcium 58, 638–648 (2015).

43. Stuttgen MC, Schwarz C. Psychophysical and neurometric detection performance under stimulus uncertainty. Nat Neurosci 11, 1091–1099 (2008).

44. Britten KH, Shadlen MN, Newsome WT, Movshon JA. The analysis of visual motion: a comparison of neuronal and psychophysical performance. J Neurosci 12, 4745–4765 (1992).

