## Supplementary figures for "Dynamic perceptual feature selectivity in primary somatosensory cortex upon reversal learning"

Ronan Chéreau et al.

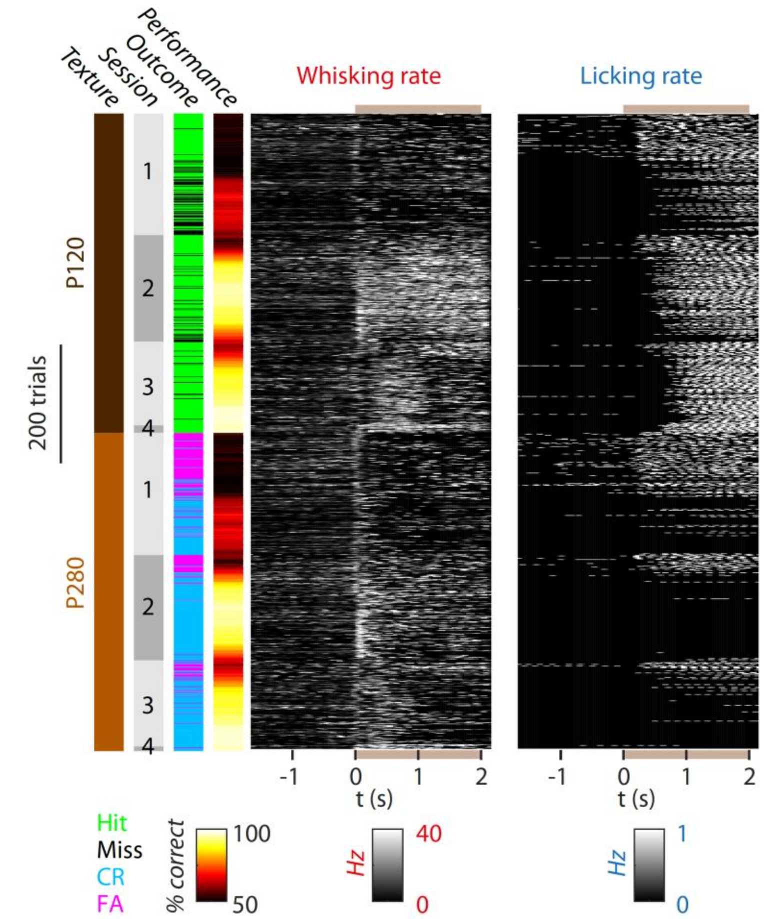


**Supplementary figure 1. Evolution of whisking and licking strategies across learning.** Single trial whisking and licking rates for a mouse during training, aligned to texture presentation onset (t=0s). Trials were sorted by texture (P120, Go stimulus and P280, No-go stimulus). For each trial, the corresponding session number, trial outcome and performance are shown. The shaded brown area indicates the time during which the texture is presented (0-2 s).

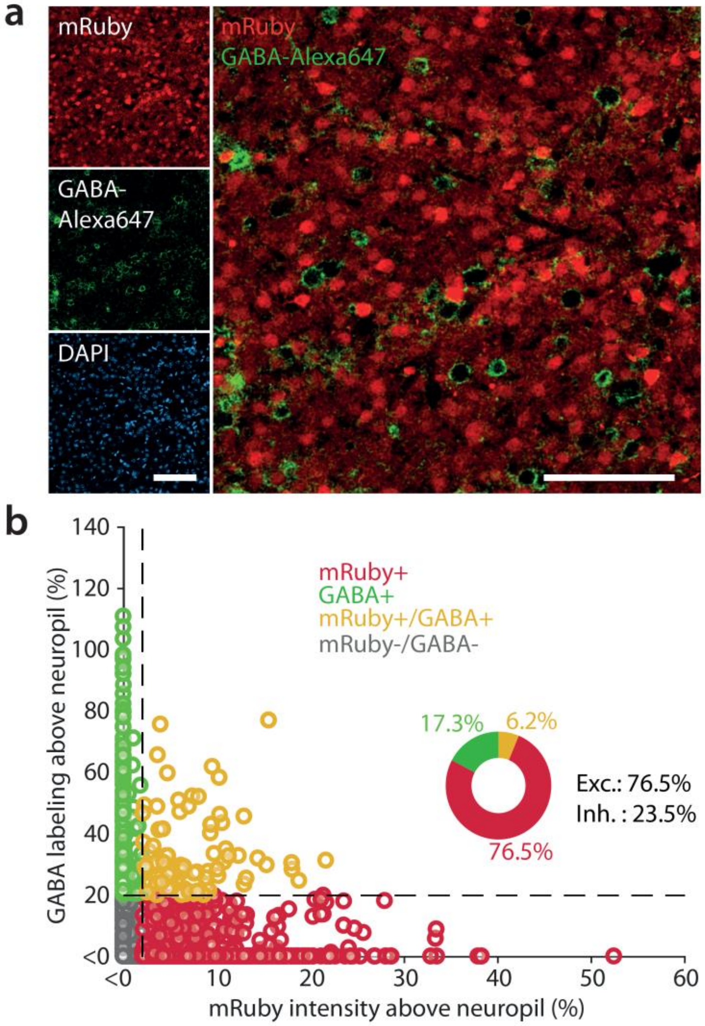


**Supplementary figure 2. Fraction of mRuby2-P2A-GCaMP6s-expressing neurons producing GABA. a**- Example of mRuby2-expression (red), GABA immunostaining (green), and DAPI labeling (blue) of L2/3 neurons in S1 on the left and a merge of the mRuby2 expression and GABA immunostaining. Note that the vast majority of GABA-positive neurons is negative for GABA and vice versa. Scale bars: 100 $\mu$m. **b**- Scatter plot comparing mRuby2 fluorescence intensity compared to GABA labeling fluorescence intensity both normalized to neuropil (each data point represents a neuron, *N*=2276 neurons measured from 5 mice). Normalized signal intensity difference above 20% was considered as labeled. Pie chart shows the percentage of neurons that were mRuby2 positive only (red), GABA positive only (green), and mRuby2+GABA positive (yellow). Only 6.2% of mRuby2-positive neurons were GABA positive, indicating that the vast majority of mRuby2/GCaMP6s-expressing neurons are pyramidal neurons.

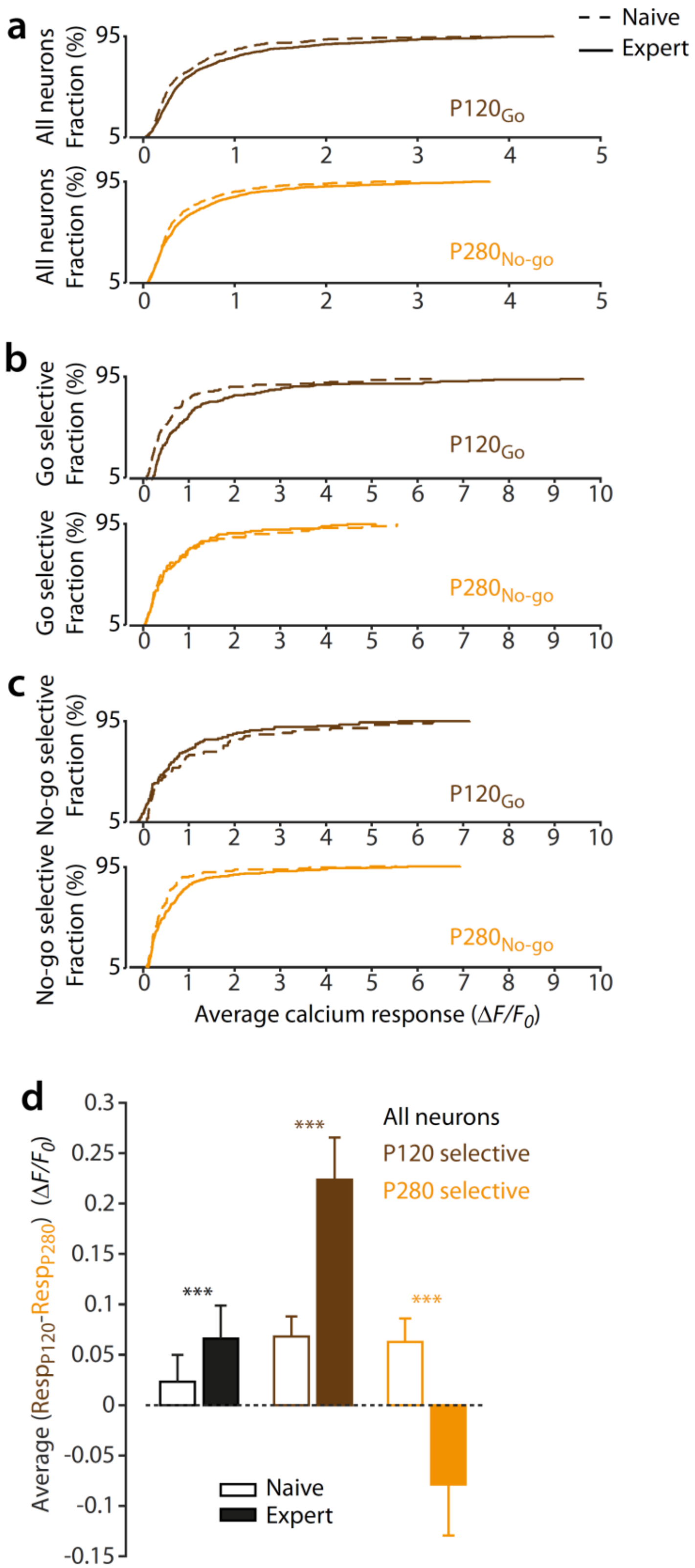


**Supplementary figure 3. Evolution of calcium responses across learning.** **a-** Cumulative distributions of the mean calcium responses per neuron for the P120 texture (Go stimulus) and

P280 texture (No-go stimulus) in naïve (dashed line) and expert sessions (continuous line). For presentation purposes, the top and bottom 5% of the population measurements are not shown. **b**- Same for P120-selective neurons only. **c**- Same for P280-selective neurons only. **d**- Average difference between normalized calcium responses of the P120 and P280 textures for all neurons (black), P120-selective neurons (brown), P280-selective neurons (orange) in naïve and expert sessions (all neurons: $N$=600 neurons in naïve and $N$=931 neurons in expert, Wilcoxon rank sum test naïve vs. expert, ***$P$=9e-5; P120-selective neurons: $N$=116 neurons in naïve and $N$=193 neurons in expert, Wilcoxon rank sum test naïve vs. expert, ***$P$=5.8e-10; P280-selective neurons: $N$=71 neurons in naïve and $N$=98 neurons in expert, Wilcoxon rank sum test naïve vs. expert, ***$P$=3.6e-4).

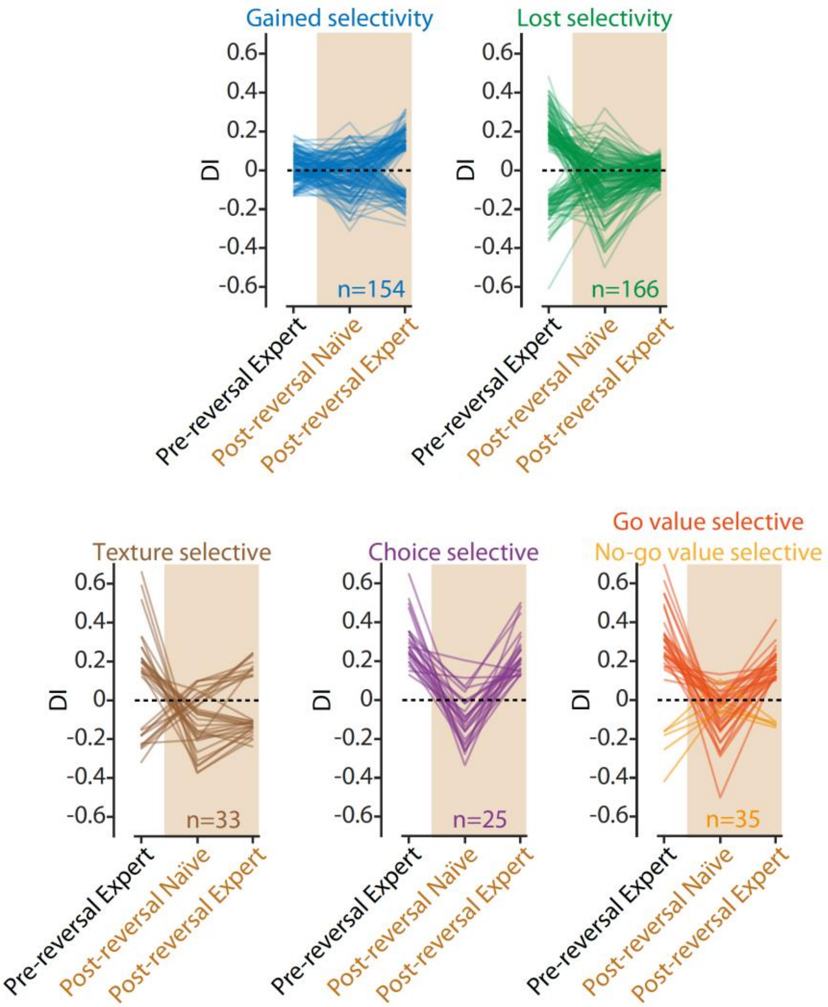


**Supplementary figure 4. Performance pre and post-reversal learning and selectivity of the neuronal classes.** Evolution of DI of all neuronal classes in the corresponding pre-reversal expert, post-reversal naïve and post-reversal expert sessions.

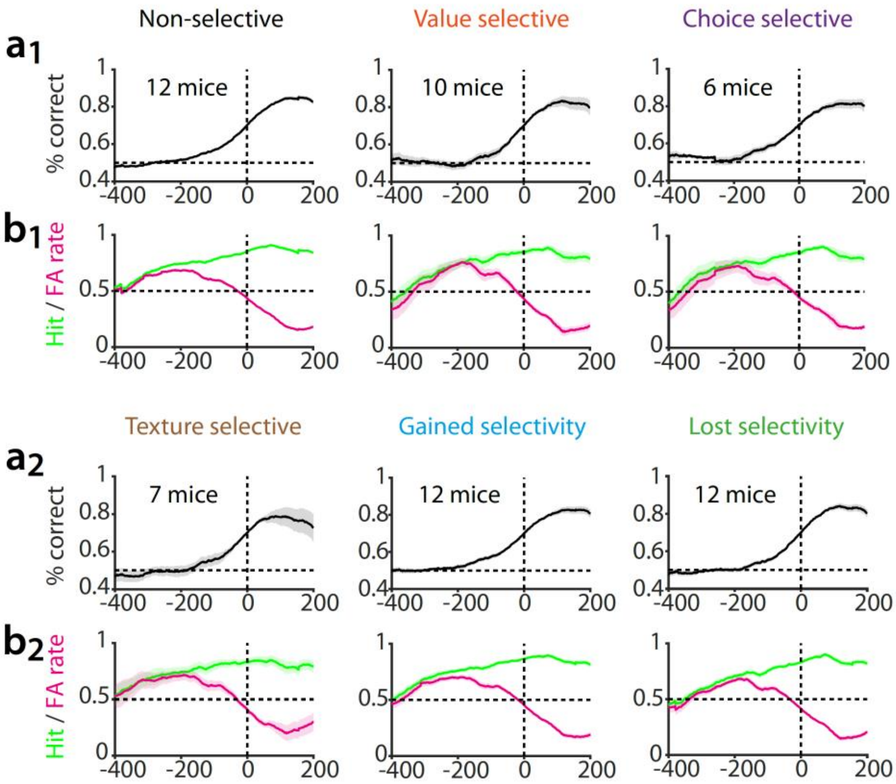


**Supplementary figure 5. Performance realignments across all identified neuronal classes during reversal learning. a**- All identified classes of neurons were not systematically detected in all mice. The average performance during reversal learning was calculated for the group of mice from which at least one neuron belonged to the given class. Performance curves were realigned to expert criterion (Non-selective neurons, *N*=412, 12 mice; Value neurons, *N*=35, 10 mice; Choice neurons, *N*=24, 6 mice; Texture neurons *N*=33, 7 mice, Neurons gaining selectivity after reversal *N*=144, 12 mice; Neurons losing selectivity after reversal *N*=152, 12 mice). **b**- Corresponding 'Hit' and 'FA' rates.

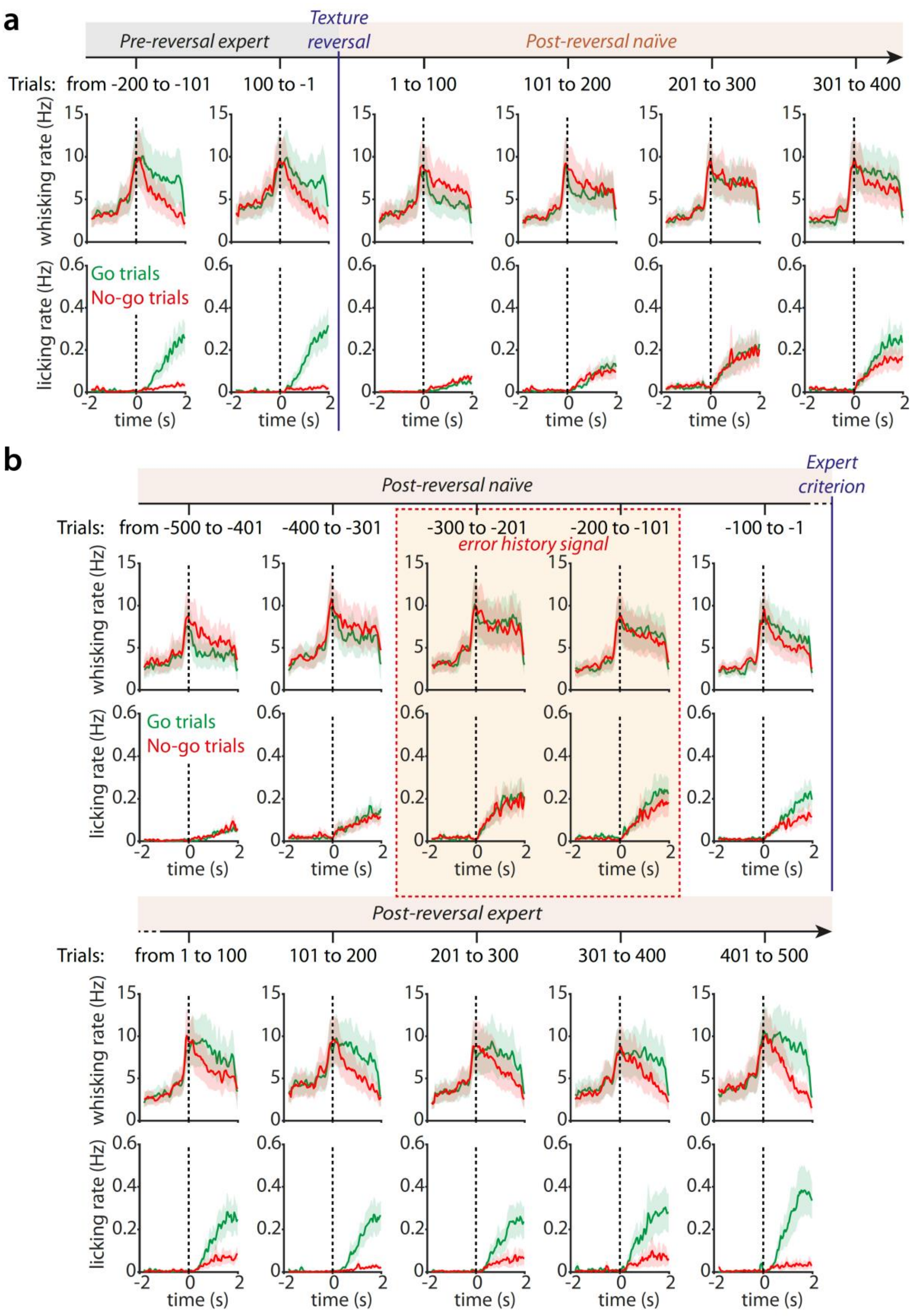
a
Pre-reversal expert
Texture reversal
Post-reversal naïve
Trials: from -200 to -101
100 to -1
1 to 100
101 to 200
201 to 300
301 to 400
whisking rate (Hz)
licking rate (Hz)
Go trials
No-go trials
time (s)
b
Post-reversal naïve
Expert criterion
Trials: from -500 to -401
-400 to -301
-300 to -201
-200 to -101
-100 to -1
error history signal
whisking rate (Hz)
licking rate (Hz)
Go trials
No-go trials
time (s)
Post-reversal expert
Trials: from 1 to 100
101 to 200
201 to 300
301 to 400
401 to 500
time (s)

**Supplementary figure 6. Whisking and licking rates pre and post texture reversal. a**- Time course of whisking (top) and licking (bottom) rates aligned to texture presentation onset (dotted line) across two intervals of 100 trials before texture reversal (P120 texture was presented on the Go trials in green, P280 on the No-go trials in red) and the first four intervals after reversal (P280 texture was presented on the Go trials in green, P120 on the No-go trials in red). We observed a decrease in whisking and licking rates for the Go trials immediately after texture reversal. This may reflect a loss of commitment to the task due a decline in the number of water rewards due to increased FAs and Misses. Whisking and licking rates gradually increase 200 trials after texture reversal as mice start to randomly lick again upon presentation of the Go and No-go stimulus. **b**- Average time course of whisking (top) and licking (bottom) rates aligned to texture presentation (dotted line) from five bins of 100 trials before and after the expert criterion (70% performance). Within the -300 to -101 interval (dashed red area), from which an error history signal was detected in the value selective neurons, whisking and licking strategies only moderately change. When mice became experts, the whisking rates remained elevated upon the Go stimulus presentations, which may reflect whisker movements induced by the motor action of licking. Shaded areas represent s.e.m.

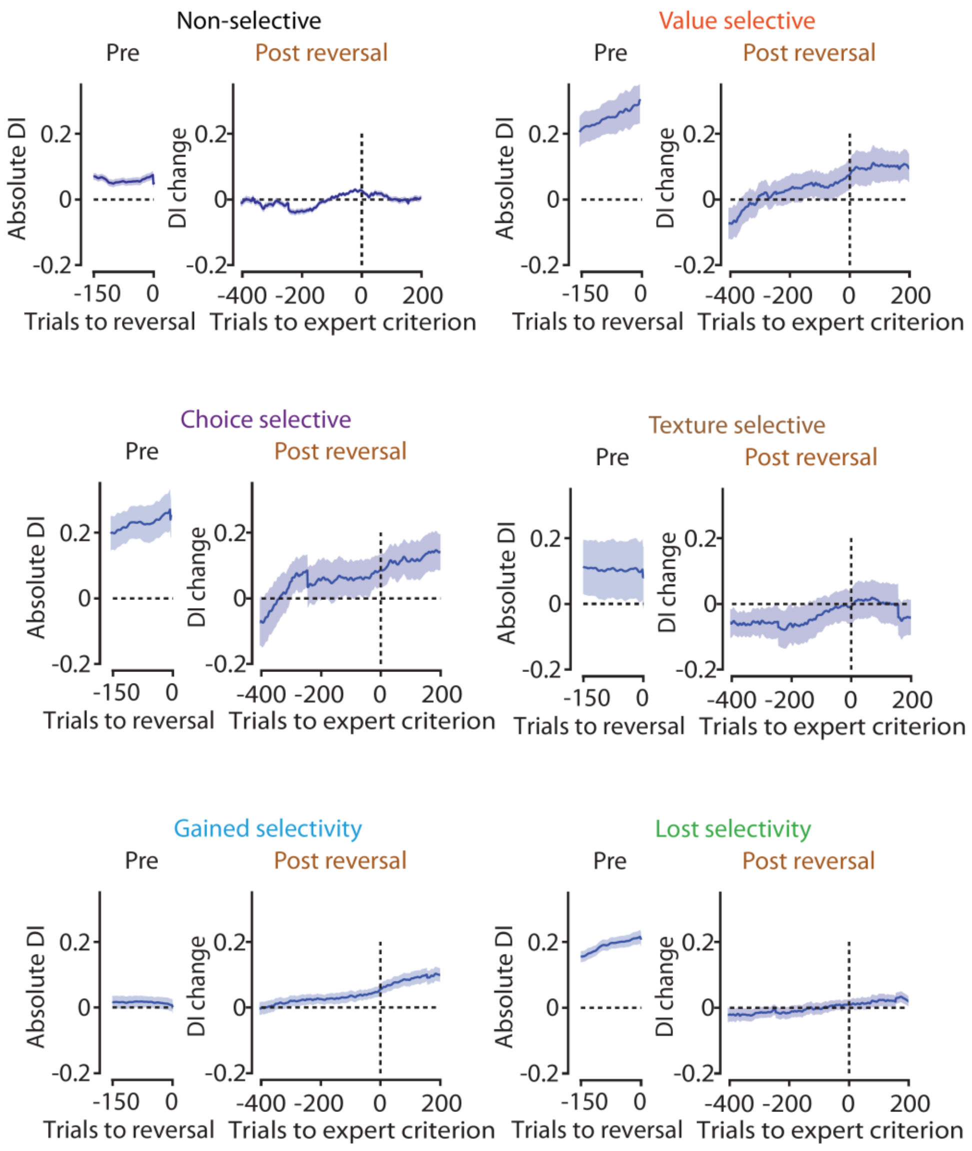


**Supplementary figure 7. Evolution of selectivity for all identified neuronal classes.** DI profiles for the various classes of neurons. For each panel, the graph on the left shows the average absolute DI across mice over the last 150 trials pre-reversal. The right graph shows the average change in DI from the absolute DI on the left, post-reversal, and realigned across mice to the expert criterion (Non-selective neurons, *N*=412, 12 mice; Value neurons, *N*=35, 10 mice; Choice neurons, *N*=24, 6 mice; Texture neurons *N*=33, 7 mice, Neurons gaining selectivity after reversal *N*=144, 12 mice; Neurons losing selectivity after reversal *N*=152, 12 mice).
